# Crystal structure of human PACRG in complex with MEIG1

**DOI:** 10.1101/783373

**Authors:** Nimra Khan, Dylan Pelletier, Simon Veyron, Nathalie Croteau, Muneyoshi Ichikawa, Corbin Black, Ahmad Abdelzaher Zaki Khalifa, Sami Chaaban, Igor Kurinov, Gary Brouhard, Khanh Huy Bui, Jean-François Trempe

## Abstract

In human, the Parkin Co-Regulated Gene (PACRG) shares a bidirectional promoter with Parkin, a gene involved in Parkinson’s disease, mitochondrial quality control and inflammation. The PACRG protein is essential to the formation of the inner junction between doublet microtubules of the axoneme, a structure found in flagella and cilia. PACRG interacts with tubulin as well as the meiosis expressed gene 1 (MEIG1) protein, which is essential for spermiogenesis in mice. However, the 3D structure of human PACRG is unknown. Here, we report the crystal structure of the C-terminal domain of human PACRG in complex with MEIG1 at 2.1 Å resolution. PACRG adopts an *α*-helical structure with a loop insertion that mediates a conserved network of interactions with MEIG1. Using the cryo-electron tomography structure of the axonemal doublet microtubule from the flagellated protozoan *Chlamydomonas reinhardtii*, we generate a model of a mammalian microtubule doublet inner junction, which reveals how PACRG interacts with tubulin subunits in both the A- and B-tubules. Furthermore, the model shows that MEIG1 interacts with *β*-tubulin on the outer surface of the B-tubule, facing towards the central pair of the axoneme. We also model the PACRG-like protein (PACRGL), a homolog of PACRG with potential roles in microtubule remodelling and axonemal inner junction formation. Finally, we explore the evolution of the PACRG and Parkin head-to-head gene structure and analyze the tissue distribution of their transcripts. Our work establishes a framework to assess the function of the PACRG family of proteins and its adaptor proteins in the function of motile and non-motile cilia.

## Introduction

Loss-of-function mutations in Parkin cause early-onset recessive Parkinson’s disease (PD) in humans [1]. Parkin is an E3 ubiquitin ligase that ubiquitinates proteins selectively on damaged mitochondria following activation by PINK1, a mitochondrial ubiquitin kinase (reviewed in [2]). Parkin and PINK1 are conserved across nearly all metazoans and appear to play a key role in mitochondrial quality control. Indeed, deletion of the *parkin* or *pink1* genes in *Drosophila* results in mitochondrial defects that cause muscle degeneration [3–5]. While Parkin or PINK1 knockout (KO) mice do not exhibit significant muscle or neuronal degeneration on their own [6, 7], mitochondria-linked stress and infection-induced inflammation can induce neurodegeneration in those strains [8–10]. Parkin KO mice are more susceptible to intracellular bacterial pathogens [11], and missense mutations in Parkin are associated with an increased susceptibility to a type-1 reaction in leprosy, a disease caused by a chronic *Mycobacterium leprae* infection [12]. Furthermore, a previous study showed that susceptibility to leprosy was strongly associated with genetic variants in the promoter region of Parkin [13]. This promoter is shared with the Parkin Co-Regulated Gene (PACRG), which is linked head-to-head with Parkin on the opposite DNA strand, with only 204 bp separating the two transcripts [14]. While the function of Parkin has been intensively studied over the last 20 years, far less is known about PACRG. Furthermore, the significance of the co-regulation of these two genes remains unclear.

PACRG encodes a 29 kDa protein that is conserved in all metazoans, as well as flagellated protozoans [15]. Immunostaining shows that Lewy bodies, which are intracellular aggregates of *α*-synuclein found in the brain of PD patients, contain PACRG [16, 17]. However, the normal physiological role of PACRG is in the formation of the axoneme, a microtubule doublet structure found in motile and non-motile cilia and flagella. Early proteomics studies showed that cilia axonemes isolated from human epithelial bronchial cells contain PACRG [18]. Loss of PACRG causes male sterility in the *quaking^viable^* (*qk^v^*) mouse, a recessive deletion of the *Parkin*, *PACRG* and *QK* genes with a severe neurological phenotype [15, 19]. Mouse PACRG is highly abundant in testis and is required for spermiogenesis [15]. Loss of PACRG results in hydrocephalus in both PACRG KO and *qk^v^* mice [19, 20]; PACRG levels are indeed elevated in ependymal cells lining the cerebral ventricles, suggesting that PACRG is required for cilia motility. Knockdown of the two PACRG paralogues in trypanosome, a protozoan parasite, results in paralysis of the flagellum and loss of outer doublet microtubules [21]. Consistent with its role in stabilizing the axoneme outer-doublet, recombinant human PACRG co-sediments with microtubules and can bundle microtubules in vitro, thus showing that it binds directly to tubulin [22]. The PACRG protein localizes to the axoneme in trypanosome as well as in *Chlamydomonas reinhardtii*, a flagellated green alga [23]. Recently, electron microscopy of *C. reinhardtii* axoneme doublet microtubules showed that PACRG localizes at the inner junction of the two tubules, in alternance with FAP20 [24]. In addition to its role in the formation of motile cilia and flagella, PACRG also localizes to a subset of non-motile cilia in sensory neurons from the nematode *Caenorhabditis elegans*, where it regulates signaling processes linked to gustatory plasticity [25]. However, PACRG is not required for ciliogenesis *per se*, nor for intraflagellar transport. Thus, PACRG is critical for the formation of *functional* motile and non-motile cilia.

In addition to its interaction with tubulin, PACRG also binds to the meiosis expressed gene 1 product (MEIG1), a 11 kDa protein that is required for spermiogenesis [26]. MEIG1 KO mice display male sterility, but don’t have other phenotypic traits found in PACRG KO mice, suggesting a more specific role in sperm maturation. PACRG and MEIG1 colocalize to the manchette of elongating spermatids, and both are required for the manchette localization of the cargo protein SPAG16, a sperm central apparatus protein [27]. Furthermore, the manchette localization of MEIG1 requires PACRG, whereas the localization of PACRG is unaffected by the loss of MEIG1. PACRG is thus upstream of MEIG1, with the latter playing a more specific role in sperm-specific cargo recruitment. However, MEIG1 also appears to stabilize PACRG expression in cells. The NMR structure solution of MEIG1 revealed a small compact domain with four exposed aromatic side-chains that are required for binding and stabilization of PACRG in bacterial cells [28]. Thus, PACRG and MEIG1 appear to form a stable heterodimer complex, but the structure of PACRG and how it binds MEIG1 are unknown.

Here, we report the crystal structure of the C-terminal domain of PACRG bound to MEIG1. While the N-terminal 69 amino acids of PACRG are disordered, PACRG^70-257^ folds into a set of helical repeats similar to those found in HEAT repeats and VHS domains. MEIG1 interacts primarily with a long loop located on the N-terminal end of the domain. Homology modelling using the high-resolution cryo-electron microscopy structure of the axoneme doublet microtubule reveals that MEIG1 likely interacts with the B-tubule and acts as an adaptor for other proteins that bind at the doublet microtubule inner junction. We identify rare human variants of PACRG that disrupt the PACRG:MEIG1 complex and predict that the PACRG-like protein likely does not bind MEIG1 nor the inner junction protein FAP20. Finally, we explore the evolution of the Parkin-PACRG head-to-head gene structure in metazoans and discuss the implications for their functions.

## Results

### Crystal structure of the human PACRG:MEIG1 complex at 2.1 Å resolution

Since previous reports indicated that mouse PACRG was stabilized by MEIG1 in *E. coli* [28], we co-expressed the two human orthologues from a single plasmid with a cleavable hexahistidine (His_6_) tag on the N-terminus of MEIG1 for affinity purification. We confirmed that human PACRG is indeed stabilized by co-expression with MEIG1 (Suppl. Figure S1). PACRG co-purifies with MEIG1 and the two proteins co-elute on size-exclusion chromatography at ∼40 kDa, consistent with the predicted molecular weight of the heterodimer (Suppl. Figure S2). However, the complex did not produce crystals. To facilitate crystallization, we removed the N-terminal 69 amino acids of PACRG, which are predicted to be disordered and are poorly conserved in other PACRG orthologues (Suppl. Figure S3). However, PACRG^Δ1-69^ did not co-purify with His_6_-MEIG1 following co-expression. We had previously noted that glutathione-S-transferase (GST)-PACRG^Δ1-69^ forms insoluble inclusion bodies. We exploited that system to produce large amounts of the GST-fusion protein, which was resolubilized in urea and then diluted and dialyzed to refold the GST domain [29]. The partially refolded fusion protein was then incubated with purified His_6_-MEIG1, and the complex was affinity purified. The affinity tags were cleaved and the products resolved by size-exclusion chromatography, which confirmed that the two proteins form a heterodimer complex of ∼33 kDa (Figure 1A).

**Figure 1.**
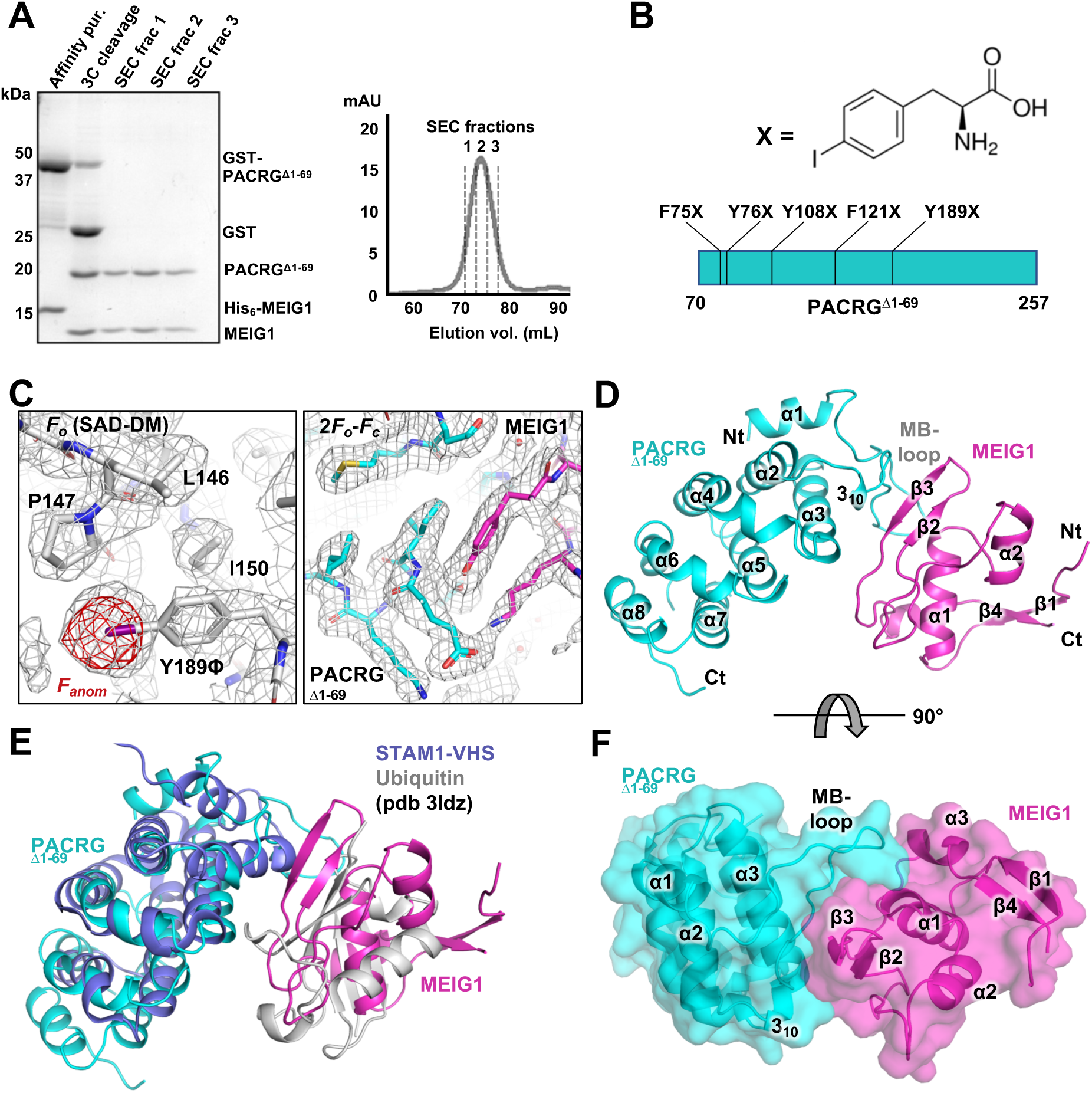
Crystal structure of the PACRG:MEIG1 complex. **(A)** Purification of PACRG^Δ1-69^: MEIG1 by GST-affinity and size-exclusion chromatography (SEC). Left, SDS-PAGE stained with Coomassie Blue showing the different steps of purification. Lanes of the SEC fractions (profile on the right) show formation of a complex between MEIG1 and PACRG^Δ1-69^. **(B)** Chemical structure of *p*-iodo-*L*-phenylalanine and sites of incorporation in PACRG. **(C)** Left, electron density maps after SAD phasing and density modification (grey mesh), with the final refined model of the Y189ϕ derivative (sticks). The anomalous difference density map (red mesh) shows a strong signal at the iodine position in Y189ϕ. Right, refined 2*F_o_-F_c_* electron density maps (mesh) for the native complex of PACRG^Δ1-69^ (cyan) and MEIG1 (magenta). **(D)** Cartoon representation of the native complex of PACRG^Δ1-69^ (cyan) and MEIG1 (magenta). Secondary structure elements are labeled and numbered. **(E)** Superposition of the STAM1-VHS domain bound to ubiquitin (violet and grey) on the PACRG^Δ1-69^:MEIG1 complex. **(F)** Surface representation of PACRG^Δ1-69^:MEIG1, rotated at 90° from the view in panel D, to highlight the interaction between the MB-loop in PACRG and a groove in MEIG1.

The MEIG1:PACRG^Δ1-69^ complex formed crystals that diffracted to 2.1 Å resolution in a tetragonal space group (Table 1 and Suppl. Figure S4). However, the structure could not be solved by molecular replacement using the mouse MEIG1 solution structure or homology models of PACRG. Selenomethionine (SeMet)-labeled PACRG^Δ1-69^ did not express, and the anomalous signal from SeMet-MEIG1:PACRG^Δ1-69^ crystals was insufficient to solve the structure. Thus, we produced *p*-iodo-*L*-phenylalanine derivatives of PACRG^Δ1-69^ using site-specific incorporation at the *amber* TAG codon to obtain anomalous signal from iodine [30, 31]. We selected five aromatic residues in human PACRG^Δ1-69^ for site-specific incorporation, with either low sequence conservation or potential surface exposure to alleviate loss of interaction with MEIG1 or disruption of the fold by the bulkier side-chain (Figure 1B and Suppl. Figure S3). All five mutants expressed and formed complexes with MEIG1, and mass spectrometry confirmed incorporation of a single *p*-iodo-*L*-phenylalanine (ϕ) residue in all of them (Suppl. Figure S5). Three mutants produced diffracting crystals (F75ϕ, Y108ϕ, and Y189ϕ), and the Y189ϕ mutant produced the best diffraction for phasing by single-wavelength anomalous dispersion (Figure 1C and Table 1). The resulting mutant structure was then used as a search model for molecular replacement of the native data set. The native and mutant structures are very similar (0.15 Å backbone rmsd), confirming that *p*-iodo-*L*-phenylalanine does not disrupt the fold of PACRG. Finally, to confirm that the structure obtained was not an artefact from the refolding procedure, we obtained crystals by limited proteolysis with subtilisin of co-expressed full-length PACRG:MEIG1. These crystals were in the same space group as MEIG1:PACRG^Δ1-69^ and diffracted to 2.6 Å (Table 1). The structure is nearly identical to MEIG1:PACRG^Δ1-69^ (0.15 Å backbone rmsd) and no additional residues were observed upstream of Thr70, which is consistent with the N-terminus being disordered.

**Table 1.** X-ray crystallography data collection and refinement statistics.

|  | **Native**<br>**MEIG1:PACRG<sup>Δ1-69</sup>** | **iodo-Y189Φ**<br>**MEIG1:PACRG<sup>Δ1-69</sup>** | **Proteolysis**<br>**MEIG1:PACRG** |
| --- | --- | --- | --- |
| **Wavelength (Å)** | 0.9793 | 1.4760 | 0.9795 |
| **Resolution range (Å)** | 47.6 - 2.1 (2.175 - 2.1) | 47.64 - 2.09 (2.16-2.09) | 47.3 - 2.6 (2.72 - 2.6) |
| **Space group** | P4<sub>1</sub>2<sub>1</sub>2 | P4<sub>1</sub>2<sub>1</sub>2 | P4<sub>1</sub>2<sub>1</sub>2 |
| **Unit cell dimensions (Å)** | 67.30 67.30 158.26 | 67.37 67.37 159.72 | 66.85 66.85 158.92 |
| **Unit cell angles (deg)** | 90 90 90 | 90 90 90 | 90 90 90 |
| **Total reflections** | 138796 (14193) | 253522 (9944) | 148977 (17761) |
| **Unique reflections** | 21980 (2139) | 22406 (1769) | 11731 (1395) |
| **Multiplicity** | 6.3 (6.6) | 11.3 (5.6) | 12.6 (12.7) |
| **Completeness (%)** | 99.79 (99.95) | 99.8 (98.4) | 100 (100) |
| **Mean I/sigma(I)** | 20.25 (1.93) | 13.7 (1.1) | 9.1 (1.1) |
| **Wilson B-factor** | 47.87 | 43.83 | 54.77 |
| **R-merge** | 0.050 (1.15) | 0.093 (1.11) | 0.274 (2.87) |
| **CC<sub>1/2</sub>** | 0.999 (0.644) | 0.998 (0.733) | 0.996 (0.557) |
| **Reflections used for R-free** | 1113 (5.1%) | 2049 (9.1%) | 634 (5.4%) |
| **R-work** | 0.190 (0.280) | 0.229 (0.359) | 0.214 (0.300) |
| **R-free** | 0.227 (0.323) | 0.249 (0.340) | 0.284 (0.363) |
| **# non-hydrogen atoms** | 2166 | 2156 | 2112 |
| **macromolecules** | 2083 | 2076 | 2066 |
| **ligands** | 6 | 6 | 25 |
| **water** | 77 | 89 | 21 |
| **Protein residues** | 254 | 254 | 253 |
| **RMS (bonds, Å)** | 0.008 | 0.009 | 0.007 |
| **RMS (angles, deg)** | 1.08 | 0.953 | 0.883 |
| **Ramachandran favored (%)** | 96 | 98.4 | 97.2 |
| **Ramachandran allowed (%)** | 4 | 1.6 | 2.8 |
| **Ramachandran outliers** | none | none | none |
| **Clashscore** | 1.96 | 3.67 | 6.57 |
| **Average B-factor** | 60.1 | 52.0 | 58.9 |
| **macromolecules** | 60.1 | 52.1 | 58.9 |
| **ligands** | 93.9 | 51.6 | 59.6 |
| **solvent** | 58.4 | 48.9 | 58.8 |

Statistics for the highest-resolution shell are shown in parentheses.

The structure of PACRG^Δ1-69^ in the complex consists of a core of 7 *α*-helices that fold into parallel repeats, preceded by an N-terminal helix and a 3_10_ helix (Figure 1D). Continuous electron density is observed in PACRG from a.a. 70 to 255, except for a.a. 203-219, a disordered loop located between *α*6 and *α*7 at the C-terminus. Intriguingly, a longer isoform of PACRG resulting from an alternative splicing event results in a longer insertion in that same loop (Suppl. Figure S6). Fold analysis with the PDBeFold server reveals similarity between PACRG and HEAT repeat proteins such as VHS domains and the RNA polymerase II-binding domain of Pcf11 [32]. The overall structure of the complex is notably similar to the structure of the STAM1 VHS domain bound to ubiquitin, where MEIG1 occupies a position similar to ubiquitin (Figure 1E). This is consistent with this type of fold being involved primarily in protein-protein interactions. However, co-expression with ubiquitin did not allow co-purification of PACRG, suggesting that PACRG does not bind ubiquitin.

The structure of human MEIG1 is similar to the NMR structure of mouse MEIG1 (1.34 Å backbone rmsd) and shows electron density for a.a. 5 to 88. MEIG1 interacts with a long loop in PACRG (which we name the MB-loop) located between helices *α*1 and *α*2, as well as residues in the *α*3 helix (Figure 1F). The buried surface area between the two proteins (694 Å^2^) is relatively small but involves 12 hydrogen bonds and 6 salt bridges (Figure 2A), which likely account for the strength of the interaction. Contacts notably involve the MEIG1 residues Trp50, Lys57, Phe66 and Tyr68, which were shown to be essential for binding PACRG [28]. To determine which amino acids in PACRG are essential for binding to MEIG1, we performed site-directed mutagenesis in full-length PACRG and co-expressed with His_6_-MEIG1. PACRG mutants that don’t bind should not co-purify with His_6_-MEIG1 following immobilized metal affinity chromatography. The results show that Lys93, Glu99, Ile100, Glu101, His132, Glu136, and His137 are required for the interaction (Figure 2B and Suppl. Figure S7). These residues are conserved in all animals that express MEIG1, which comprise all vertebrates as well as some invertebrates such as *Aplysia*, but not insects nor single-celled organisms (Figure 2C). Furthermore, mutation of His106, which makes a crystal contact with a MEIG1 chain from another asymmetric unit, does not affect the interaction, confirming that the reported heterodimer is the one that mediates the interaction in solution. Overall, these results confirm that PACRG forms a stable complex with MEIG1 as observed in the crystal structure.

**Figure 2.**
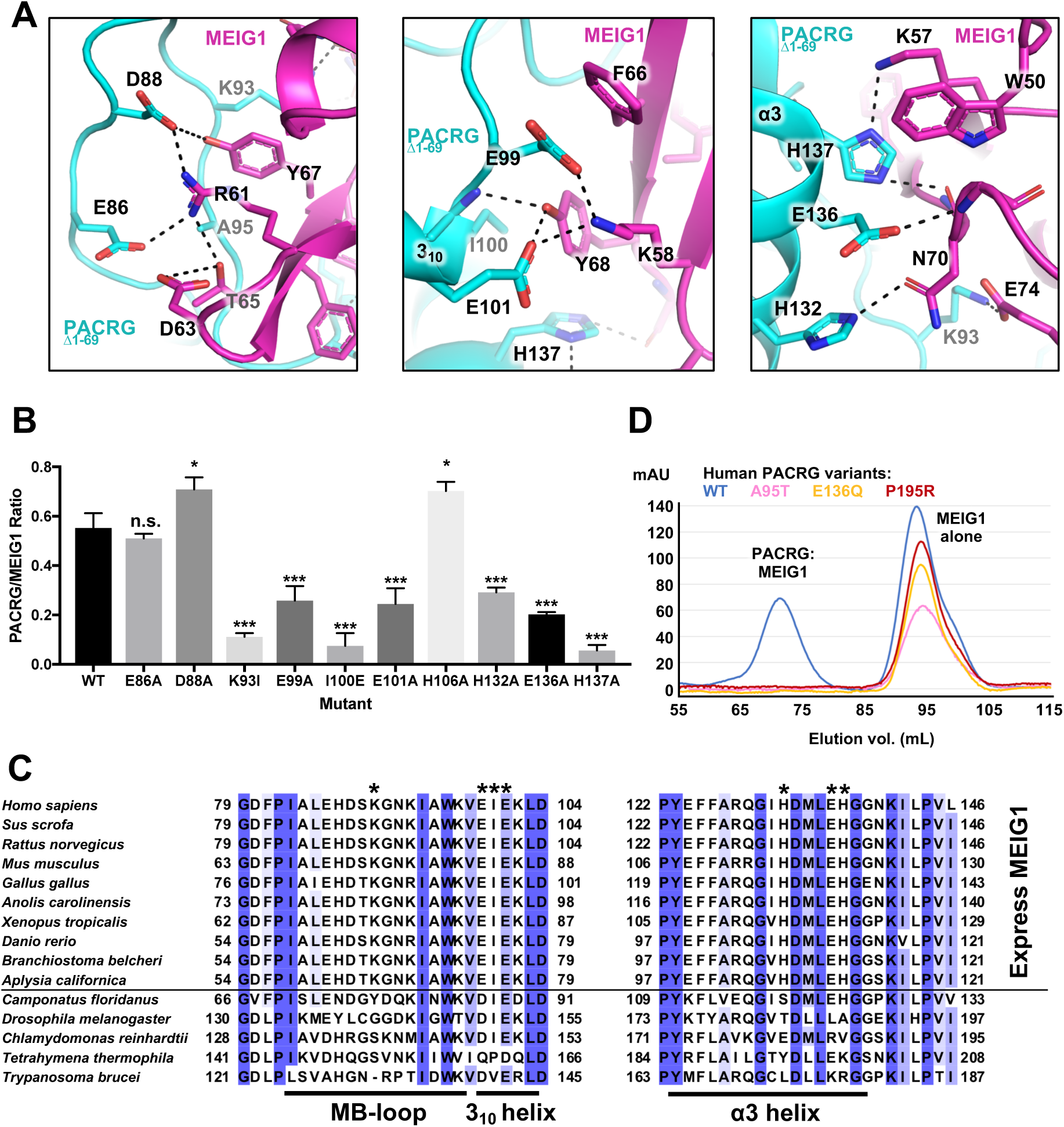
The PACRG:MEIG1 interface reveals a conserved network of polar interactions. **(A)** Close-up views and interactions observed in the crystal structure of the native PACRG^Δ1-69^:MEIG1 complex. Dashed lines indicate hydrogen bonds. **(B)** Interaction assay for the binding of PACRG mutants to His_6_-MEIG1. The two proteins were co-expressed and purified by immobilized metal affinity chromatography (IMAC). Products were analyzed by SDS-PAGE followed by Coomassie Blue staining. The ratio of PACRG to His_6_-MEIG1 was calculated by densitometry (*n*=4). One-way ANOVA and Dunnet’s tests were performed for comparison with the wild-type (WT) interaction (* for *α* ≤0.05, *** for *α* ≤0.001). **(C)** Sequence alignment of PACRG orthologues across different organisms. Species above the horizontal line also express MEIG1. The asterisks indicate residues in human PACRG that are required for binding MEIG1. Figure generated with *Jalview*. Blue shading indicates degree of conservation. **(D)** Size-exclusion chromatography of WT and rare human variants of PACRG, following co-expression with His_6_-MEIG1 and IMAC. The WT shows two peaks for the complex (at 73 mL) and His6-MEIG1 alone (at 94 mL). No complex is observed for the mutants.

### Model of PACRG:MEIG1 bound to the inner junction of the axonemal doublet microtubule

Electron microscopy studies showed that PACRG localizes outside the A-tubule of the axonemal doublet and positions itself at the inner junction of the two tubules [22, 24]. However, it is unclear how PACRG interacts with tubulin and how MEIG1 positions itself within the axonemal doublet. In an accompanying manuscript, the structure of the axonemal doublet microtubule from *C. reinhardtii* was solved by cryo-electron tomography to a resolution of 3.6 Å [33]. High-resolution electron density was observed at the inner junction, which allowed model building of PACRG, FAP20 and a number of other axonemal proteins. The structure revealed that *C. reinhardtii* PACRG (CrPACRG) and FAP20 alternate at the inner junction of the A and B-tubules (Figure 3A). The CrPACRG C-terminal helical domain, corresponding to human (Hs)PACRG^Δ1-69^, interacts simultaneously with the doublet microtubule A1 and B10 protofilaments and makes contacts with both *α*- and *β*-tubulin (Figure 3B,C). Electron density was observed for the N-terminus of CrPACRG, which folds into an *α*-helix that binds *β*-tubulin on the B-tubule. This helix is then followed by a long segment that adopts a “triple-helix”-like conformation that interacts with *β*-tubulin subunits in protofilaments A1 and A13.

**Figure 3.**
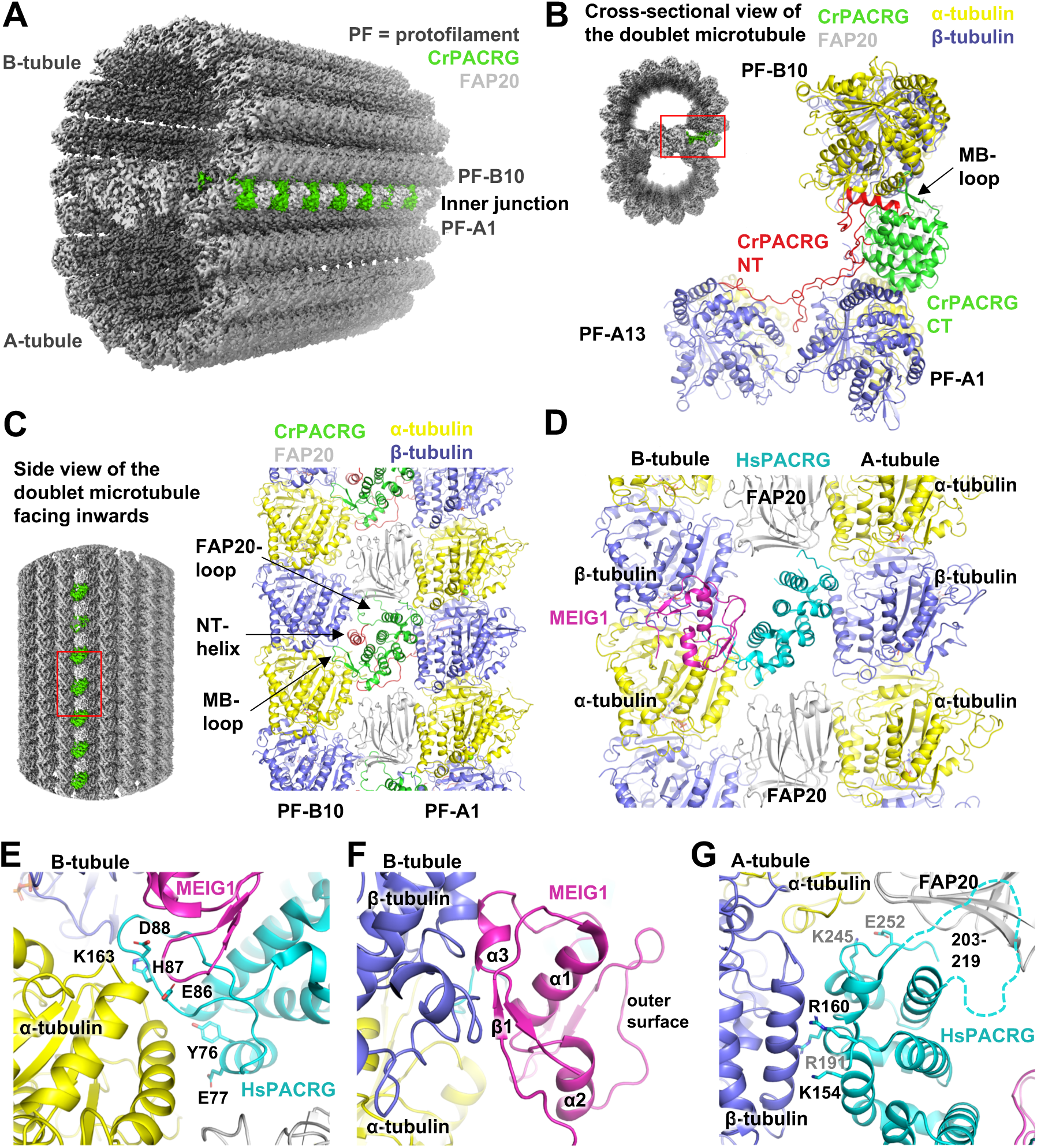
Structural model of PACRG:MEIG1 in the axonemal doublet microtubule. **(A)** *Chlamydomonas reinhardtii* axonemal doublet microtubule structure solved by cryo-electron tomography [33]. PACRG and FAP20 are colored at the inner junction between the A- and B-tubules. Tubulin protofilaments (PF) are in gray. **(B)** Cross-sectional view of the axonemal doublet showing the interaction of PACRG with the A1 and B10 PFs. The N-terminus (NT, red) of PACRG makes extensive interactions with tubulin. The loop that corresponds to the MEIG1-binding loop in HsPACRG is on the outer surface of the inner junction. **(C)** Side-view of the axonemal doublet viewed from the outside facing inwards, as viewed from the central pair. **(D)** Side-view of the homology model (facing inwards) of the inner junction of the *human* axonemal doublet. HsPACRG (cyan) interacts with *α*-tubulin on the B-tubule, where MEIG1 (magenta) would also be located. The C-terminus of HsPACRG interacts with both *α*- and *β*-tubulin on the A-tubule. FAP20 is found on both sides of PACRG. **(E-G)** Close-up views of the interactions mediated by HsPACRG:MEIG1 in the homology model of the inner junction. Critical residues in PACRG that are in contact with tubulin are labeled. The dashed line in panel G indicate the position of the disordered loop (a.a. 203-219) in HsPACRG, which is ordered and binds FAP20 in the *Chlamydomonas* structure.

In order to understand how the human PACRG:MEIG1 complex would interact with the doublet microtubule, we have built a homology model by superposing our crystal structure on the CrPACRG coordinates in the cryo-EM structure [34]. HsPACRG^70-257^ and CrPACRG^119-307^ are very similar (0.71 Å backbone rmsd) and thus the two proteins likely mediate very similar types of interactions with tubulin (Figure 3D). The MB-loop is located on the B-tubule side and interacts primarily with *α*-tubulin. Notably, we find that residues Glu86 and Asp88, which interact with MEIG1 but whose mutations do not affect MEIG1 binding (see Figure 2), are in very close proximity with a basic patch in *α*-tubulin comprising Lys163 and Lys164 (Figure 3E). This positions MEIG1 very close to *β*-tubulin, with potential interactions between the *α*3 helix of MEIG1 and the H3 helix of *β*-tubulin (Figure 3F). This would place MEIG1 on the outer surface of the inner junction of the doublet microtubule. HsPACRG would also interact with the A-tubule, notably via three invariant basic residues (Lys154, Arg160, Arg191) that would bind to an acidic patch in *β*-tubulin (Figure 3G). Two highly conserved residues also interact with *α*-tubulin from the A-tubule (Lys245, Glu252). Finally, the CrPACRG structure shows that the loop connecting *α*6 and *α*7 (a.a. 203-219), which is disordered in the HsPACRG crystal structure, interacts with FAP20.

To test the microtubule remodeling activity of PACRG:MEIG1, we performed a co-polymerization reaction with tubulin purified from cow brain. Polymerized microtubules were then visualized using negative stain electron microscopy (Figure 4). Addition of wild-type (WT) PACRG:MEIG1 to the reaction resulted in the formation of protofilament sheet structures and fewer closed microtubules. The PACRG:MEIG1 complex is thus a microtubule remodelling protein that can stabilize atypical lattice structures such as those found at the inner junction of doublet microtubules.

**Figure 4.**
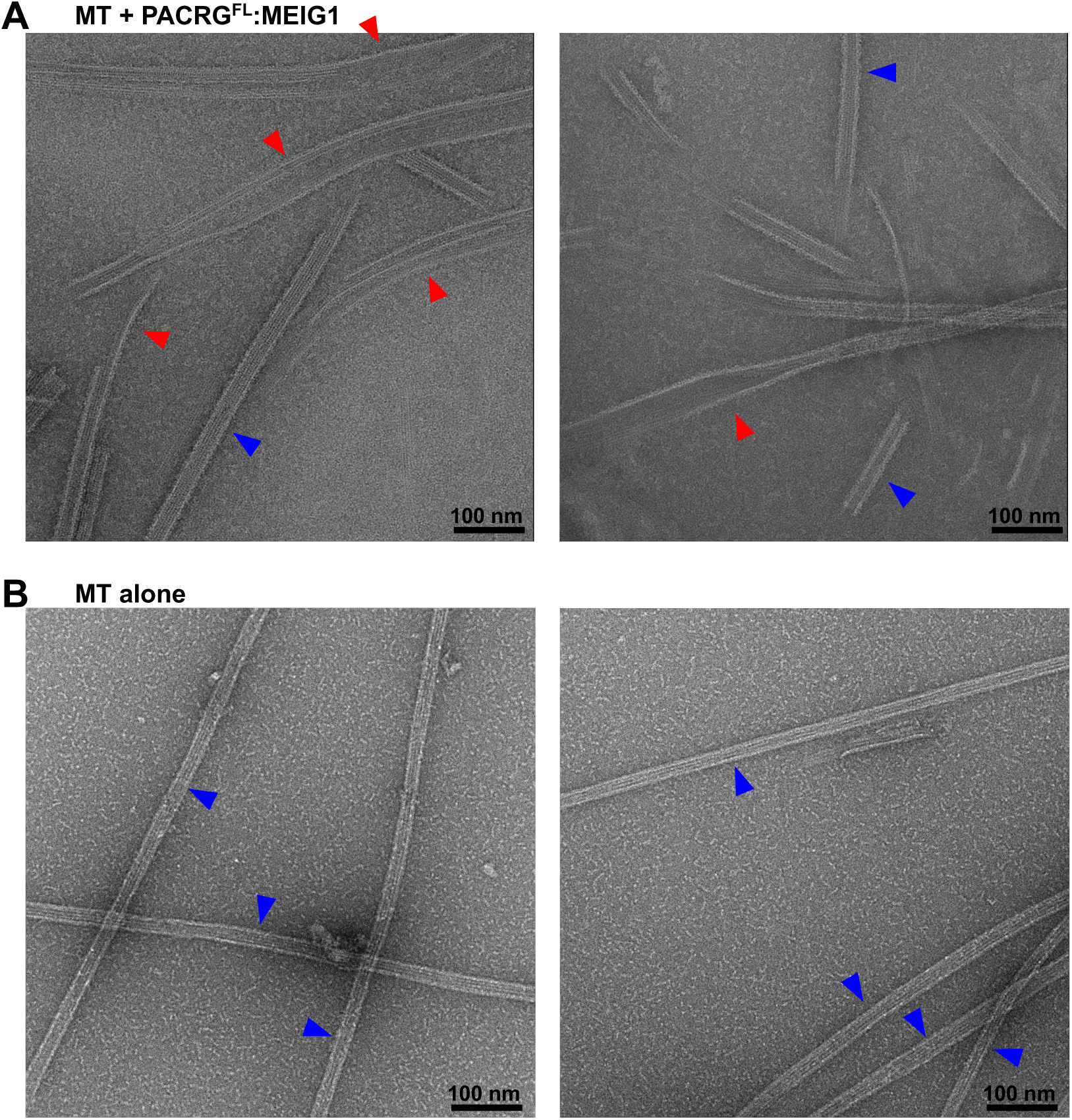
PACRG:MEIG1 stabilizes atypical microtubule lattice structure. **(A)** Negative stain electron microscopy images of microtubules (MT), co-polymerized in the presence of PACRG^FL^:MEIG1. Red arrows indicate sheet-like structure of polymerized tubulin, indicating that PACRG stabilizes another lattice structure. Blue arrows indicate typical MTs. **(B)** Negative control showing typical MT structures obtained by polymerization in the absence of PACRG:MEIG1.

### Rare human PACRG genetic variants impair MEIG1 binding

In order to explore the impact of the PACRG interactions in human health, we searched the GnomAD database for missense variants in the protein at residues involved in binding MEIG1 or tubulin [35]. By excluding variants with only one allele, we identified 7 rare variants with allele frequencies ranging between 1 and 10 per 100,000 individuals (Table 2). We introduced these mutations in the full-length PACRG:MEIG1 complex for expression and purification. Three of those missense mutations failed to form a complex with MEIG1 on gel filtration (Figure 2D). Two of those missense mutations, A95T and E136Q, are in proximity to MEIG1 in the crystal structure and likely disrupt this interaction (Figure 2A). The P195R mutation is located at the interface of *β*-tubulin in the B-tubule, but the mutation likely disrupts the *α*6 helix and thus unfolds PACRG. Thus, the A95T, E136Q and P195R rare variants disrupt the function of PACRG:MEIG1, which may explain their low frequency in the human population.

**Table 2.** Human PACRG variants that affect MEIG1 or tubulin interactions

| **Human PACRG variants** | **Allele count** | **Total alleles** | **Allele frequency** | **Role of residue** |
| --- | --- | --- | --- | --- |
| Y76C | 8 | 282714 | 2.8E-05 | Binding to α-tubulin (B-tubule) |
| E86K | 2 | 251328 | 8.0E-06 | Binding to α-tubulin (B-tubule) and MEIG1 |
| A95T | 26 | 251066 | 1.0E-04 | Binding to MEIG1 |
| E136Q | 2 | 282826 | 7.1E-06 | Binding to MEIG1 |
| R160Q | 10 | 282322 | 3.5E-05 | Binding to β-tubulin (A-tubule) |
| P195R | 8 | 282736 | 2.8E-05 | Binding to β-tubulin (A-tubule) |
| E252K | 10 | 282336 | 3.5E-05 | Binding to α-tubulin (A-tubule) |

Data obtained from the GnomAD database

The remaining four variants are potentially involved in making interactions that stabilize the inner junction of the axonemal doublet. Notably, the variants Y76C and E86K may affect the interaction of PACRG with *α*-tubulin in the B-tubule (Figure 3E), and the variants R160Q and E252K would affect interaction with the A-tubule (Figure 3G).

### Modelling of the PACRG-like protein suggests a role in microtubule remodelling independent of FAP20 and MEIG1

Sequence homology search with PACRG identifies the PACRG-like protein (PACRGL), a poorly characterized protein with 23% sequence identity with PACRG (Figure 5A). Using the crystal structure of PACRG, we have created a homology model of PACRGL (Figure 5B). The model shows that PACRGL has a very short loop between *α*6 and *α*7, strongly suggesting that it cannot interact with FAP20. Furthermore, five residues in PACRG that are essential for MEIG1 binding (Lys93, Ile100, His132, Glu136 and His137) are not conserved in PACRGL, suggesting that it does not bind MEIG1 (Figure 5C). Residues at the predicted tubulin interfaces are not entirely conserved, although it is still unclear what the molecular determinants are for these interactions. Still we find that the tubulin-interacting PACRG residues Tyr76, His87, Lys154, Arg191, and Lys245 are conserved in PACRGL (Tyr84, His95, Lys164, Lys198, and Lys237, respectively). Since the two proteins likely have the same fold, the implication is that PACRGL may indeed bind tubulin, but not FAP20, and thus would play a different role than PACRG in microtubule remodelling.

**Figure 5.**
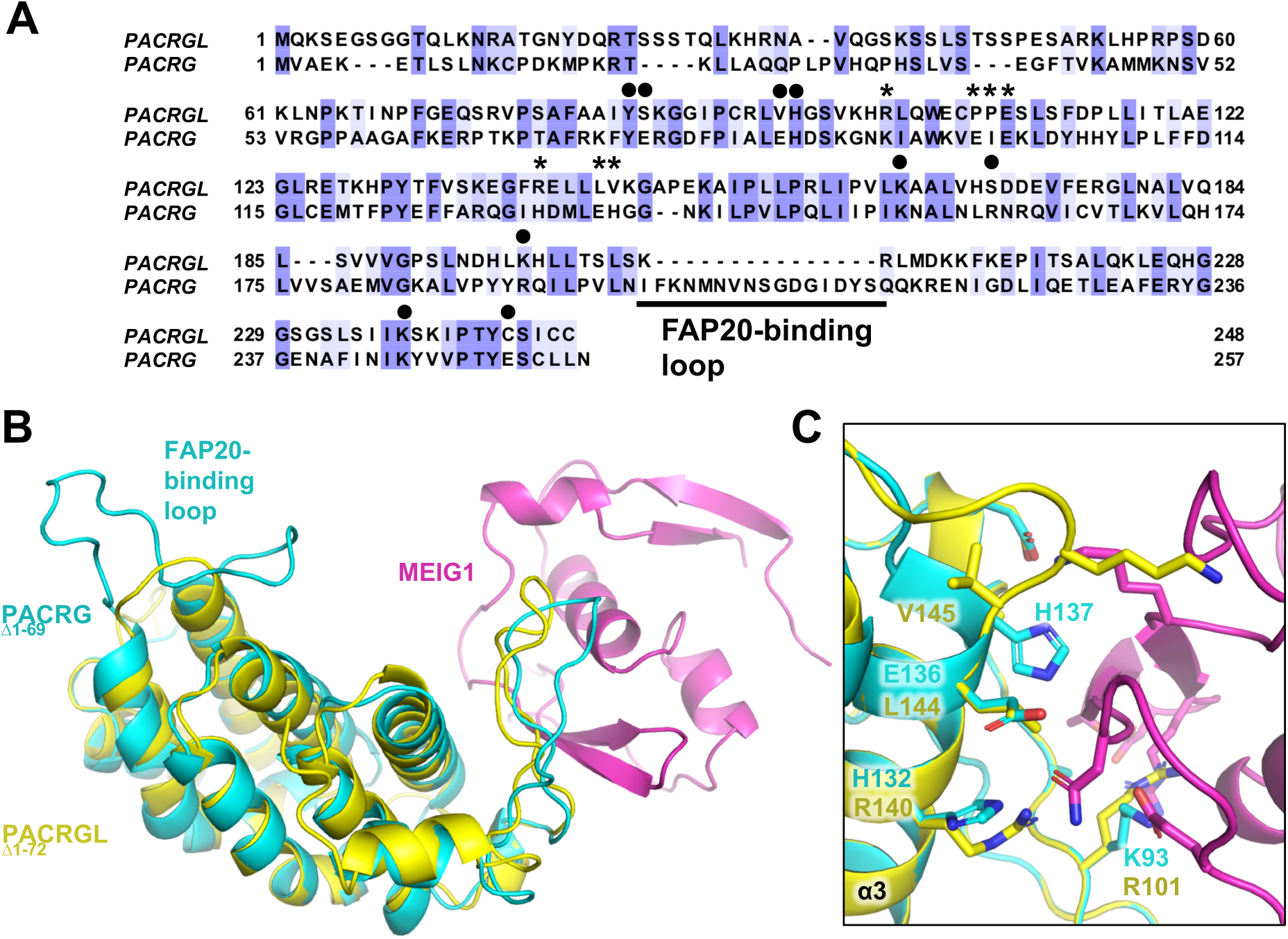
Homology model of the PACRG-like protein suggest a role in microtubule remodelling independent of FAP20 and MEIG1. **(A)** Sequence alignment of the human PACRG-like protein (PACRGL) with human PACRG. Blue shading indicates degree of conservation. Asterisks indicate residues important for binding MEIG1. Black circles highlight residues that contact tubulin in the axonemal doublet tubule structure. **(B)** Homology model of PACRGL^Δ1-72^ (yellow) superposed on the crystal structure of PACRG^Δ1-69^:MEIG1 (cyan and magenta). **(C)** Close-up view of the PACRG:MEIG1 interface, showing the side-chain of residues critical for the interaction. The side-chain of corresponding PACRGL residues are also shown.

### Conservation of the PACRG-Parkin head-to-head gene structure and transcription regulation

Gene pairs such as Parkin and PACRG, which share bi-directional promoters, may also share a common function and have partially overlapping expression patterns [14]. Furthermore, their relative orientation is not random in mammals, with the head-to-head orientation being preferred for gene duos separated by less than 600 bp [36]. To establish whether Parkin and PACRG function in a common biological pathway, we have analyzed the conservation of the PACRG and Parkin gene structure. In all amniotes (terrestrial animals), PACRG and Parkin are found in a head-to-head orientation, with most species having less than 600 bp between the two genes, except in a few mammals and birds (Table 3). The relationship breaks in amphibians, which do not have a Parkin gene. Fishes and invertebrates have Parkin and PACRG, but they are typically located on different chromosomes, except for the elephant shark. This is consistent with head-to-head gene duos starting to accumulate in tetrapods [36]. Finally, it is worth noting that single-celled organisms do not have Parkin, but have PACRG if they are ciliated or flagellated (Table 3). PACRG is thus tightly associated with organisms that have flagella or motile cilia based on the axonemal doublet microtubule structure.

**Table 3.** Evolution of PACRG-Parkin head-to-head (H2H) gene structure and conservation of PACRG and MEIG1

| Species name (latin) | Common name | PACRG-Parkin H2H? | Space (kb) | %ID PACRG | %ID MEIG1 | Comments |
| --- | --- | --- | --- | --- | --- | --- |
| *Homo sapiens* | Human | Yes | 0 | 100.0 | 100.0 |  |
| *Pan troglodytes* | Chimp | Yes | 0 | 100.0 | 97.7 |  |
| *Gorilla Gorilla* | Gorilla | Yes | 0 | 100.0 | 96.6 |  |
| *Sus scrofa* | Pig | Yes | 240 | 96.9 | 94.3 |  |
| *Rattus norvegicus* | Rat | Yes | 0 | 96.1 | 92.1 |  |
| *Mus musculus* | Mouse | Yes | 0 | 92.1 | 87.5 |  |
| *Ursus maritimus* | Polar bear | Yes | 200 | 97.3 | 95.5 |  |
| *Myotis lucifugus* | Little brown bat | Yes | 0 | 90.0 | 90.9 |  |
| *Gallus gallus* | Chicken | Yes | 0 | 70.4 | 77.0 |  |
| *Calypste anna* | Hummingbird | Yes | 150 | 85.0 | 80.7 |  |
| *Anolis carolinensis* | Green anole | Yes | 0 | 92.2 | 77.0 |  |
| *Alligator mississippiensis* | American alligator | Yes | 0 | 81.4 | 81.8 |  |
| *Terrapene mexicana triunguis* | Three-toed box turtle | Yes | 0 | 82.9 | 87.5 |  |
| *Xenopus laevis* | African clawed frog | n.a. | n.a. | 90.0 | 75.0 | No parkin/PINK1 |
| *Xenopus tropicalis* | Tropical clawed frog | n.a. | n.a. | 91.3 | 75.0 | No parkin/PINK1 |
| *Danio rerio* | Zebrafish | No | n.a. | 89.6 | 71.6 |  |
| *Callorhynchus milii* | Elephant shark | Yes | 0 | 76.8 | 79.5 |  |
| *Salmo salar* | Atlantic salmon | No | n.a. | 88.7 | 75.3 |  |
| *Aplysia californica* | Sea slug | No | n.a. | 82.3 | 61.0 |  |
| *Drosophila melanogaster* | Fruit fly | No | n.a. | 59.9 | none | No MEIG1 |
| *Caenorhabditis elegans* | Nematode worm | No | n.a. | 33.5 | none | No MEIG1 |
| *Chlamydomonas reinhardtii* | Green algae | n.a. | n.a. | 63.7 | none | No Parkin/PINK1, MEIG1 |
| *Tetrahymena thermophila* | Ciliate | n.a. | n.a. | 60.5 | none | No Parkin/PINK1, MEIG1 |
| *Paramecium tetraurelia* | Ciliate | n.a. | n.a. | 58.7 | none | No Parkin/PINK1, MEIG1 |
| *Trypanosoma brucei* | Kinetoplastids | n.a. | n.a. | 57.4 | none | No Parkin/PINK1, MEIG1 |
| *Oxyrrhis marina* | Dinoflagellate | n.a. | n.a. | None | 38.9 | No Parkin/PINK1, PACRG |

Given that the head-to-head structure of the PACRG-Parkin genes is conserved in amniotes, we sought to determine whether their expression was co-regulated as a result of acting in related pathways. We therefore analyzed their mRNA expression profiles across different tissues using the *Protein Atlas* consensus dataset [37]. We extended our analysis to MEIG1, PACRGL, and other axonemal proteins (SPAG16, FAP20, and FAP206), as well as PINK1 and DJ-1, two proteins implicated in PD and which regulate mitochondrial homeostasis [38]. Furthermore, we also included LRRK2, a kinase associated with both leprosy and PD [12]. We found a strong correlation (*r* = 0.88) between the expression of Parkin and PINK1, with both expressing at very high levels in mitochondria-rich tissues such as muscles and the central nervous system (Figure 6). Likewise, we found a significant correlation between PACRG and MEIG1 (*r* = 0.74), with high expression in reproductive tissues such as testis and Fallopian tubes as well as some brain areas. PACRG expression is notably very high in the pituitary gland, which contains multiciliated cells [39, 40]. PACRG and MEIG1 levels both correlate well with other axonemal proteins. However, there was little correlation between PACRG and Parkin (*r* = 0.18), with some overlap in the basal ganglia and cerebral cortex. LRRK2 expression did not correlate with Parkin nor with PACRG, although the two tissues with the highest levels of LRRK2, namely lung and granulocytes, show significant levels of PACRG. On the other hand, the expression of PACRGL correlates with PACRG (*r* = 0.58), MEIG1 (*r* = 0.69), and other axonemal proteins, suggesting that PACRGL is also an axonemal protein.

**Figure 6.**
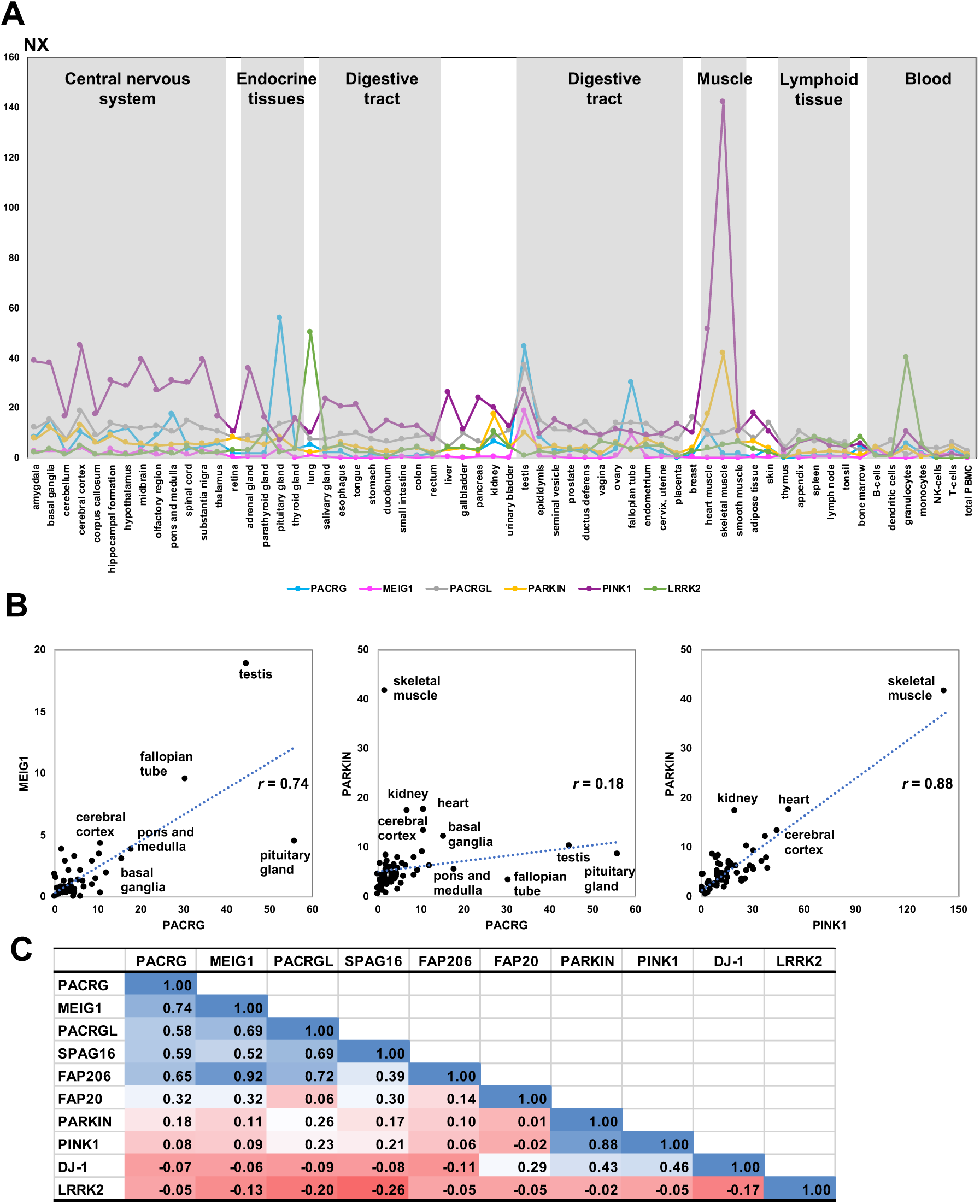
Transcriptomics analysis of PACRG and other genes across human tissues. Data retrieved from the *Protein Atlas* consensus dataset for mRNA expression in different human tissues and blood cells [37] **(A)**. Consensus Normalized eXpression (NX) levels for each gene, sorted by tissue type. **(B)** Correlation graphs of NX levels for three selected pairs of genes (PACRG/MEIG1, PACRG/PARKIN, PARKIN/PINK1). Tissues showing high levels of expression of one or both genes are labeled. Correlation coefficients (*r*) are indicated. **(C)** Table of pairwise correlation coefficients between tissue mRNA expression levels, color-coded from low (red) to high (blue) values.

## Discussion

In this study, we report the high-resolution crystal structure of human MEIG1 bound to PACRG with a N-terminal deletion. The structure was solved using incorporation of the non-natural amino acid *p*-iodo-*L*-phenylalanine, with the iodine atom providing strong anomalous signal that we used for experimental phasing (Figure 1). The structure revealed a heterodimer complex with a high number of polar interactions. PACRG adopts a helical repeat fold similar to that found in HEAT proteins and interacts with MEIG1 via an atypical loop that we named the MB-loop. The interaction was validated by mutagenesis and revealed residues in PACRG that are essential for binding MEIG1 (Figure 2). These residues are conserved in all organisms that also express MEIG1, which comprises all vertebrates as well as invertebrates that express motile sperm cells such as mollusks. However, other metazoans as well as ciliated protozoans do not have MEIG1 but yet have PACRG, hinting to the latter playing a more fundamental role in axonemal doublet formation, whereas MEIG1 evolved later to have an adaptor function.

The structure of the *Chlamydomonas* axonemal doublet by cryo-electron tomography by Bui and colleagues [33] enabled us to build a homology model of human PACRG:MEIG1 at the inner junction of an axonemal doublet microtubule (Figure 3). The *Chlamydomonas* doublet structure notably reveals that PACRG and FAP20 alternate at the inner junction, with each protein interacting with two protofilaments, on the A- and B-tubules. Consistent with this prediction, we find that addition of PACRG:MEIG1 to a tubulin polymerization assay interferes with single microtubule formation and results in the formation of protofilament sheets, as PACRG binds to one side of the protofilament and stabilize an atypical lattice structure (Figure 4). Strikingly, homology modeling also shows that MEIG1 would interact with the A-tubule, on the external side of the inner junction. In the 9+2 axoneme structure, this would position MEIG1 towards the interior of the axoneme, facing the central doublet. According to Li et al. [27], MEIG1 would act as an adaptor protein for cargo protein SPAG16, which is itself interacting with central pair proteins [41]. Thus, the position of MEIG1 in the axonemal doublet is consistent with its proposed role as an adaptor protein for cargo located in the center of the 9+2 axoneme.

While the significance of our findings on human health is still uncertain, the PACRG:MEIG1 structure enables us to predict how naturally-occurring variants in PACRG might affect its function. We have discovered three PACRG variants (A95T, E136Q, and P195R) that disrupt the interaction with MEIG1 (Figure 2D). Furthermore, we have identified four additional variants (Y76C, E86K, R160Q, and E252K) that may disrupt interactions with tubulin protofilaments at the inner junction the axonemal doublet (Table 2 and Figure 3). We were able to purify complexes formed by these four mutants, thus confirming that these mutations do not impair the interaction with MEIG1. Notably, while PACRG-Glu86 forms a salt bridge with MEIG1-Arg61, mutations E86A and E86K do not disrupt the complex (Figure 2B,D). However, this salt bridge contacts *α*-tubulin in the A-tubule of the homology model (Figure 3E), and thus the E86K variant would perturb a network of interaction that might be essential for maintaining the axonemal inner junction. While the phenotype of humans carrying these variants is unknown, we would predict that they display phenotypes similar to those found in ciliopathies [42]. Loss-of-function mutations in PACRG will likely impair fertility and might also lead to neurodevelopmental and respiratory defects. Indeed, mice lacking PACRG display male sterility and hydrocephalus [19, 20]. Furthermore, PACRG is expressed in the epithelial bronchial cells, where cilia play a critical role in the clearance of mucus [18]. Thus, patients with loss-of-function mutations in PACRG may also display respiratory defects such as chronic bronchitis and sinusitis. We find that PACRG and MEIG1 levels are elevated in the Fallopian tubes, where motile cilia play an essential role in the maturation and transport of the oocyte [42]. PACRG/MEIG1 mutations may therefore also impair female fertility. Finally, it has not yet been established whether mammalian PACRG localizes at the primary cilia, a non-motile axoneme-based structure that coordinates a plethora of cellular signalling pathways [43]. *C. elegans* PACRG was found to localize to non-motile cilia involved in G-protein signalling and gustatory plasticity [25]. Thus, if PACRG is essential to the function of primary cilia, loss-of-function PACRG mutations would have long-ranging effects on Sonic hedgehog signalling, G-protein signalling and so on, which could have a profound impact on human health. Future studies should aim to determine the role of PACRG in the function of motile and primary cilia in mammals.

PACRG was discovered as a gene sharing a bidirectional promoter with Parkin [14]. We found that this head-to-head arrangement of the two genes is conserved in amniotes (Table 3). Yet, the significance of this observation remains unclear. Transcriptomics analysis shows that the tissue expression patterns of the two genes are not correlated, although the two genes are co-expressed in many tissues (Figure 6). In mouse, PACRG and Parkin are both expressed in testis and brain, and the single knockout of PACRG leads to an increase in Parkin levels, suggesting some cross-talk [19]. A potential route of interaction between the two genes might be in sperm maturation. Mitochondria play an important role in sperm development as they form an array with microtubules that promotes the elongation of the tail of the sperm, which eventually becomes the flagella [44]. The mature spermatid tail is filled with mitochondria, which are in direct contact with the flagellar axoneme to provide the energy required for motility. Parkin is a well-established mitochondrial quality control protein, and Parkin-null Drosophila have mitochondrial dysfunction that results in a defect of spermatid individualization at the late stage of sperm maturation [3]. Parkin and PACRG are on different chromosomes in Drosophila (Table 3); however, it is possible that their relationship was strengthen later in evolution via formation of a head-to-head gene pair. Other tissues or cells with motile cilia, such as ependymal cells, will also have high energy requirements, where coupling between mitochondria and axoneme would be essential. Thus, we speculate that the two genes are co-regulated only in a subset of tissues, under the control of a tissue-specific transcription factor that can transcribe both genes.

Another intriguing link between Parkin and PACRG is their involvement in immunity and inflammation. Early studies by Schurr and colleagues found that single-nucleotide polymorphisms (SNPs) in the PACRG-Parkin promoter region were strongly associated with leprosy [13]. Parkin and PINK1 have recently been implicated in mitochondria-dependent inflammation, in response to bacterial infection or mitochondrial stress [8–10]. Intriguingly, SNPs and missense variants in LRRK2, a gene associated with autosomal dominant PD, have also been linked to susceptibility, inflammatory responses and type-1 reactions in leprosy [12,45,46]. LRRK2 is a large protein of the ROC-COR family of protein kinases, which regulates intracellular trafficking via phosphorylation of Rab GTPases, which are key mediators of pathways implicated in PD [47]. Intriguingly, LRRK2 phosphorylates two Rab GTPases, Rab8A and Rab10, which regulate ciliogenesis [48, 49]. Pathogenic mutations in LRRK2, which increase its kinase activity, impair primary cilia formation as a result of Rab hyper-phosphorylation. In adult mice, LRRK2 is highly expressed in the lung and lymph node [50], which is consistent with the expression pattern observed in human, where it is found in the lung and granulocytes (Figure 6). While there is no overall correlation between the expression of LRRK2 and any other genes we investigated (Figure 6), PACRG is expressed in lung and granulocytes, which point to a role in immunity that warrants further investigations.

Our crystal structure of PACRG:MEIG1 enabled us to model the structure of the PACRGL protein (Figure 5). To our knowledge, there is no publication on this protein, which appears to be expressed in a broad range of tissues in human (Figure 6). It is highly expressed in sperm and its tissue distribution correlates to some degree with PACRG and MEIG1, suggesting that their functions overlap. Furthermore, PACRGL expression correlates well with SPAG16 and FAP206, which is consistent with PACRGL being an axonemal protein. On the basis of homology modelling, we predict that PACRGL will not bind FAP20, but will likely binds tubulin in a manner similar to PACRG. This would therefore make PACRGL an axonemal microtubule inner junction protein that binds at sites distinct from PACRG. Inner junction proteins play a critical role not only in 9+2 axoneme structures found in motile cilia and flagella, but also in the regulation of microtubule doublets and singlet-to-doublet transitions that occur in non-motile primary cilia [51]. However, the identity of inner junction proteins regulating primary cilia is unknown. Finally, while PACRGL has a loop that is similar in length to that of the MB-loop in PACRG, its sequence is not compatible with MEIG1-binding (Figure 5). PACRGL may bind another, yet unknown, adaptor protein that binds at the inner junction. Furthermore, we observe that PACRG and MEIG1 tissue distribution are not perfectly correlated, with PACRG being expressed in tissues with little to no MEIG1 (e.g. heart and granulocytes). Thus, it is also possible that the MB-loop in PACRG binds other adaptor proteins that enable other proteins to bind at the inner junction of various axoneme microtubule structures. While we do observe that the PACRG:MEIG1 complex broadly resembles the STAM1-VHS:ubiquitin complex (Figure 1E), PACRG lacks residues that are critical for ubiquitin binding and indeed we could not detect an interaction between ubiquitin and PACRG. It is however intriguing that proteomics study of primary cilia performed by proximity labeling identified components of the ubiquitin pathway [52]. Thus, we cannot exclude that PACRG binds a ubiquitin chain or a ubiquitin-like molecule. Future work should seek to identify adaptor proteins for PACRG and PACRGL and determine their respective roles in the regulation of motile and non-motile cilia.

## Materials and methods

### Plasmids, antibodies and reagents

The open-reading frame for the long isoform of human PACRG^1-296^ was obtained by gene synthesis following codon optimization for expression in E. coli (DNA Express Inc., Montreal). The gene was cloned directly in the pGEX-6p1 plasmid. To generate the full-length short isoform (“FL”, 257 a.a.), primers were designed to remove a.a 205-243 by PCR mutagenesis (Suppl. Figure S8). To generate PACRG^Δ1-69^, standard restriction enzyme cloning was performed using the protocol described by New England Biolabs (NEB). PACRG*Δ*_1-69_ was amplified from the GST-PACRG^FL^ plasmid with PCR primers introduced BamHI (N-terminal) and XhoI (C-terminal) restriction sites for ligation in pGEX-6p1 with the T4 DNA ligase (NEB; Suppl. Figure S9). To generate the PACRG:MEIG1 co-expression vector the same protocol was performed as above using restriction enzyme cloning. His_6_-MEIG1 was obtained by gene synthesis from GeneArt (Thermofisher) and cloned in pRSET-A (Suppl. Figure S10). To generate pRSF-Duet-PACRG-His_6_MEIG1 (Suppl. Figure S11), pRSET-His_6_-MEIG1 was digested with NcoI and XhoI and the fragment was inserted into a pRSF-Duet vector, cleaved by the same enzymes. PACRG^FL^ (pGEX-6P-1), digested with NcoI and XhoI restriction enzymes (NEB), was inserted into pRSF-Duet vector containing His_6_-MEIG1, which was digested with NcoI and SalI, which has compatible cohesive ends with XhoI. Single-point mutants in PACRG were generated by PCR mutagenesis using the QuickChange II Site Directed Mutagenesis kit (Agilent Technologies). The pDB070.iodo.5 plasmid containing the modified tRNA and tRNA synthetase for p-iodo-L-phenylalanine incorporation was obtained from Addgene (#99397) as a gift from David Liu [31]. p-iodo-L-phenylalanine was purchased from Sigma (# I8757). The PACRG mouse C-8 monoclonal antibody was from Santa Cruz Biotechnology (# sc-373851). The His-tag polyclonal rabbit antibody was from Cell Signaling Technology (# 2366P).

### Protein expression and purification

#### Purification of His_6_-MEIG1:PACRG^FL^ and His_6_-MEIG1 alone

BL21 E. coli cells transformed with the pRSF-DUET or pRSET-A plasmids were grown overnight in LB at 37°C shaking at 180 rpm with the corresponding antibiotic. The next day, the starter culture was added to 1L or more LB, grown at 37°C shaking at 180 rpm until OD_600_ reached 0.8, at which point 0.5 mM IPTG was added for induction. Protein expression was carried out for 3 hours at 25°C. Cells were harvested at 5000 rpm for 30 min in a Sorvall type GS-3 centrifuge rotor. The cell pellet was resuspended in lysis buffer (50 mM HEPES pH 7.4, 120 mM NaCl, 5 mM MgCl2,0,25 mg.mL DNase, 0.5 mg/ml of lysozyme, 2.5 mM DTT, 0.2% triton X-100, and 1 protease cocktail inhibitor: complete ULTRA Tablets from Roche diluted in 60 ml of lysis buffer). The resuspended cell pellet was then lysed using sonication with a process time of 2 minutes, 20 seconds on pulse and 40 seconds off time. The lysate was then clarified by centrifugation at 15,000 rpm in a Beckman JA-20 rotor. Co-NTA beads were obtained from stripping 1 mL of Ni-NTA resin from Qiagen in a Bio-rad gravity column with 50 mL of 0.5M EDTA (pH 8.0), followed by 20 ml of water. 50 mL of 1 M Co(II) was then added to the NTA resin to produce Co(II) charged NTA resin. 1 mL of this resin was then washed in wash buffer (50 mM HEPES pH 7.4 and 300 mM NaCl). The cleared lysate was then added directly on the resin. The resin was then washed once in 3 column volume (cv) of wash buffer, followed by 4 washes, each of 1 cv of wash buffer. The resin was then washed twice with 1 cv of wash buffer that additionally contained 20 mM imidazole. Finally, the protein was eluted with 4 cv of 300 mM imidazole wash buffer. The purified complex was concentrated to ≤5 mL and cleaved at 4 °C overnight with 20 μg of GST-3C protease per mg of protein to cleave the His_6_ tag from MEIG1. The sample was then loaded on a 120 mL HiLoad 16/600 Superdex 75 size-exclusion chromatography column equilibrated in buffer A at 1 mL/min.

#### Purification of GST-PACRG^Δ1-69^:MEIG1

GST-PACRG^Δ1-69^ was expressed from the pGEX-6p1 plasmid using the same protocol described above. The inclusion bodies protein extraction protocol used was adapted from Li et al. [29]. Inclusion body pellets were washed, 5 mL/g of pellet, with a buffer made of 50 mM HEPES, 50 mM NaCl, 2 mM DTT, 1% Triton X-100 (pH 7.0). The pellet was resuspended 3 times using 7 mL Wheaton Dounce tissue grinder (Fisher Scientific) and centrifuged at 12,000 rpm at 4°C for 30 minutes (Sorvall SS-34 centrifuge rotor). The last wash was performed with a buffer that did not contain Triton X-100. To extract the recombinant protein from the inclusion bodies, the cell pellet was resuspended in a buffer of 8 M urea, 50 mM HEPES, 50 mM NaCl, 2 mM DTT at pH of 7.0. The resuspension was incubated at 4°C with constant rotation overnight. Once the pellet was fully dissolved the mixture was centrifuged at 15,000 rpm at 4°C for 30 minutes. The supernatant was slowly added dropwise to 450 mL of refolding buffer with 2 M urea, 50 mM HEPES, 50 mM NaCl, 2 mM DTT, 1% Tween20 (pH 7.0) to initiate protein refolding and dilution of urea. The refolding mixture was incubated at 4°C with constant slow stirring for 2 days. The mixture was then centrifuged to remove any insoluble material and then dialyzed against a 4 L solution of 20 mM HEPES, 50 mM NaCl, 2 mM DTT, 1% Tween20 (pH 7.0) for 2 days at 4°C using 3,500 molecular weight cut-off dialysis tubing (Fisherbrand). The dialyzed sample was centrifuged and kept at 4 °C or frozen at −80°C for later use. 4 mg of His_6_-MEIG1 was then added dropwise to the supernatant to induce complex formation with PACRG. The solution was left stirring overnight at 4 °C. The next day, the GST-PACRG^Δ1-69^:MEIG1 complex was purified by affinity chromatography using 1.5 mL of glutathione-sepharose resin (GE Healthcare) washed in Buffer A (50 mM of HEPES, 120 mM of NaCl, 1 mM of DTT, pH 7.4). This resin was added to dialyzed PACRGΔ1-69: MEIG1 complex and incubated at 4 °C with slow rotation for 1 h. The resin was washed Buffer A and the protein was eluted in Buffer A with 20 mM glutathione (pH 8.0). The purified complex was cleaved at 4 °C overnight with 20 μg of GST-3C protease per mg of protein to cleave the GST tag from PACRG and the His_6_ tag from MEIG1, and then further purified by size-exclusion chromatography as described above, except that the column was connected to a GST-Trap (GE Healthcare) for online removal of GST and the GST-3C protease.

#### Iodo-phenylalanine incorporation

The procedure is a modified version of a previously described protocol [31]. Briefly, the pGEX6p1-PACRG^Δ1-69^ plasmid was co-transformed with the pDB070.iodo.5 plasmid into BL21 cells. Ampicillin (100 μg/mL) and chloramphenicol (25 μg/mL) were added to the overnight culture. 25 mL of the overnight culture was added to 525 mL of LB with ampicillin and chloramphenicol, cells were grown in 37°C shaker until an optical density of 0.3. 500 μL of *p*-iodo-*L*-phenylalanine in DMSO (320 mg/mL) was added to the cells along with 110 μL of anhydrotetracycline (1 mg/mL). Once the cells reached an optical density of 0.5, IPTG was added to 0.5 mM and cells were incubated in the shaker for 3 hours at 37°C. The remainder of the protocol is identical to the inclusion body protocol described above.

### Interaction assay

His_6_-MEIG1:PACRG^FL^ WT and mutant proteins were expressed in 300 mL of LB and purified by affinity chromatography as described above. 20 μl of the imidazole elution fraction was then loaded on 12.5% acrylamide SDS-PAGE gels (Figure 2B). The protein was then stained by Coomassie Brilliant Blue and visualized using Biorad Imaging System. Densitometry was performed using the ImageJ 3 software to calculate the PACRG/His_6_-MEIG1 ratio. The experiment was performed in quadruplicates for each mutant.

### Mass spectrometry

For tandem mass spectrometry of protein digests, 10 μg of samples was denatured in 6 M urea, 1 mM EDTA, 50 mM TEAB pH 8.5. For the time course of trans phosphorylation, samples were diluted to a final concentration of 1 M urea, 10 mM EDTA and 50 mM TEAB pH 8.5 to stop the reaction. Cysteine residues were then reduced with TCEP (2 mM pH 7.0) and alkylated with iodoacetamide (10 mM solution in H_2_O freshly prepared, Sigma). Protein samples were diluted to 1 M urea with 50 mM TEAB buffer and digested with 1:100 trypsin (Sigma) 2 h at 37°C. Digested peptides were purified using C18 spin columns (Thermo Scientific). Peptides were diluted in loading buffer (2% acetonitrile, 0.1% formic acid), and 0.5 μg of peptides was captured on a Waters C18 BEH 1.0/15 mm column, washed 5 min with 4% acetonitrile, followed by a 30-min 5–40% gradient of acetonitrile in 0.1% formic acid, with a flow rate of 40 μl/min. The eluate was analyzed on a Bruker Impact II Q-TOF mass spectrometer equipped with an Apollo II ion funnel ESI source. Data were acquired in positive-ion profile mode, with a capillary voltage of 4,500 V and dry nitrogen heated at 200°C. Spectra were analyzed using the DataAnalysis and Biotools softwares (Bruker) to confirm incorporation of p-iodo-L-phenylalanine.

### Crystallization

Crystallization was performed using the MEIG1:PACRG^Δ1-69^ complex concentrated 1.7 mg/mL. Two crystallization screens were used JBScreen Basic 1 (Jena Bioscience) and JCSG plus (Molecular Dimensions). Using a Formulatrix NT8 robotic drop dispenser sitting drops of 0.3 μL of protein and 0.3 μL of mother liquor were dispensed into 96-well plates. Plates were stored at 4°C. Crystals had grown overnight in conditions from both screens. The condition that formed crystals that diffracted to 2.1 Å resolution was from a condition containing 16% w/v PEG 8000, 20% v/v glycerol and 40 mM KH_2_PO_4_. Crystals were harvested with loops directly from this drop and plunged in liquid nitrogen for diffraction experiments. The Y189ϕ-derivative was concentrated to 2 mg/mL and mixed 1 μL with 1 μL of 18% w/v PEG 8000, 15% v/v glycerol and 60 mM KH_2_PO_4_. Crystals grew in one day at 4°C and were cryo-protected in the same condition with 20% glycerol. Limited proteolysis was performed on the purified PACRG^FL^:MEIG1 complex. The protein at 2 mg/mL was digested with 1:1000 subtilisin for 30 min. 1 μL of the mixture was then incubated with 1 μL 20% w/v PEG 8000, 20% v/v glycerol and 250 mM KH_2_PO_4_. Crystals grew in one day at 4°C and were frozen directly in liquid nitrogen.

### X-ray data collection and structure determination

Diffraction data for native MEIG1:PACRG^Δ1-69^ and protease-digested MEIG1:PACRG^FL^ were collected at the CMCF beamline 08ID-1 at the Canadian Light Source. Diffraction data for p-iodo-L-phenylalanine derivatives was collected at the Advanced Photon Source NECAT 24ID-E beamline. For the MEIG1:PACRG^Δ1-69^ complex, a total of 180 images were collected with an oscillation angle of 0.5° at 0.9793 Å. For the protease-digested MEIG1:PACRG^FL^ complex, a total of 360 images were collected with an oscillation angle of 0.5° at 0.9795 Å. For the Y189ϕ – derivative of MEIG1:PACRG^Δ1-69^, a total of 360 images were collected with an oscillation angle of 0.5° at 1.476 Å. Reflections were integrated with the *XDS* and scaled with *Aimless* [53]. The Y189X derivative structure was solved by single-anomalous dispersion using *Autosol* from the software package *PHENIX* 1.12 [54, 55]. The average <*d^anom^*/*σ*> was 1.08 and 0.87 at 4.5 and 3.0 Å, respectively. Following a first round of automated model building with *ARP/WARP* [56], the model was manually adjusted with *Coot* [57] and refined in *PHENIX*. Native structures were solved by molecular replacement using the Y189X derivative structure as a search model in *PHASER* [58], and refined with Coot and PHENIX.

### Microtubule co-polymerization assay and negative stain electron microscopy

Microtubules were polymerized from α- and β-tubulin isolated from cow brain, using a protocol adapted from Ashford et al. [59] and described recently [60]. All reactions were set up in PCR tubes containing 20 μM tubulin, 20 μM WT/mutant MEIG1:PACRG, 4 mM MgCl_2_, 1 mM GTP, and BRB80 buffer (80 mM PIPES, 1 mM MgCl_2_, 1 mM EGTA, pH 6.8). BSA was used as a negative control. Microtubules were polymerized in a Peltier thermal cycler at 35°C for 1 hour (MJResearch). The microtubules were stabilized with addition of 50 μM paclitaxel then all samples were diluted to a final tubulin/microtubule concentration of 2 μM. 5 μL of each sample was dispensed onto negatively glow-discharged carbon-coated 300-mesh copper grids. Samples were left to absorb for 2 min before they were blotted and washed with BRB80 supplemented with 50 μM paclitaxel. Then samples were stained using 2% uranyl acetate. Electron microscopy was performed with a FEI Tecnai G2 Spirit Twin 120 kV transmission electron microscope was used at the Facility for Electron Microscopy Research (FEMR, McGill University). Data was collected at a magnification of either 13000 X (pixel size 8.1 Å) or 49000 X (pixel size 1.7 Å). The microscope was equipped with a Gatan Ultrascan 4000 (4k × 4k) CCD Camera and a FEI CompuStage Single-Tilt Specimen Holder.

### Homology modelling

The structure of the human PACRG-like protein was built with the automated I-TASSER server [61], using the structure of human PACRG as a template (this work; PDB 6NDU). A homology model of the human axonemal doublet was built by superposing the human PACRG:MEIG1 complex onto the *Chlamydomonas* PACRG coordinates derived from cryo-electron tomography [33]. Mammalian alpha and beta tubulin models were generated by superposing the coordinates of pig tubulin monomers (PDB 1TUB) [34] on the corresponding coordinates of *Chlamydomonas* tubulin subunits.

### Transcriptomics analysis

Consensus normalized expression levels for 55 tissue types and 7 blood cell types, created by combining the data from the three transcriptomics datasets (HPA, GTEx and FANTOM5) using an internal normalization pipeline [37], were retrieved for individual genes from the *Protein Atlas* server in TSV format (www.proteinatlas.org). Pearson correlation coefficients were calculated between each set using Excel.

## Data availability

The atomic coordinates and structure factors of the Y189Φ derivative, native MEIG1:PACRG^Δ1-69^, and subtilisin-digested MEIG1:PACRG crystal structures have been deposited in the Protein Data Bank under accession codes PDB 6NEP, 6NDU, and 6UCC, respectively. Other data are available from the corresponding author upon reasonable request.

## Acknowledgments

We thank K. Winklhofer and J. Menschede for useful discussion that contributed to the development of this study. The beamline 08ID-1 at the Canadian Light Source (CLS) is supported by the Canada Foundation for Innovation (CFI), the Natural Sciences and Engineering Research Council (NSERC), the National Research Council (NRC), the Canadian Institutes of Health Research (CIHR), the Government of Saskatchewan, and the University of Saskatchewan. We express our deepest appreciation to the staff at the CLS Canadian Macromolecular Crystallography Facility, in particular P. Grochulski, M. Fodje, and S. Labiuk. The Northeastern Collaborative Access Team (NECAT) beamlines are funded by the National Institute of General Medical Sciences from the National Institutes of Health (P30 GM124165). The Eiger 16M detector on 24-ID-E beam line is funded by a NIH-ORIP HEI grant (S10OD021527). This research used resources of the Advanced Photon Source (APS), a U.S. Department of Energy (DOE) Office of Science User Facility operated for the DOE Office of Science by Argonne National Laboratory under Contract No. DE-AC02-06CH11357. We thank T.M. Schmeing and C. Fortinez for sharing their time at the APS. We thank the McGill Pharmacology SPR/MS facility (M. Hancock) and the CFI for infrastructure support. This work was supported by a Canada Research Chair grant from the CIHR to J.-F.T., as well as a grant and studentship from NSERC to J.-F.T. and D.P.

## Contributions of authors

N.K. developed the protein expression and purification protocols, purified and crystallized the native complex, designed mutants and expression for iodo-phenylalanine incorporation, and performed analysis of the human rare variants. D.P. purified and crystallized the iodo-phenylalanine mutants, performed structure-derived mutagenesis in PACRG and performed the interaction studies. S.V. froze crystals, collected X-ray data at CLS and solved/refined the Y189Φ and subtilisin structures. N.C. performed protein purification and performed crystallization and functional studies. N.K., M.I., and C.B. performed the microtubule co-polymerization and electron microscopy assay. S.C. and G.B. purified and provided tubulin. A.A.Z.K. and K.H.B. shared the cryoEM model and contributed intellectually to the development of the project. I.K. collected the data at APS on the Y189Φ derivative. J.-F.T. froze crystals, collected X-ray data and solved/refined the native complex structure, performed homology modelling and transcriptomics analysis, wrote the manuscript and assembled the figures.

