## Supplemental data for "Crystal structure of human PACRG in complex with MEIG1"

Running title: Crystal structure of human PACRG:MEIG1

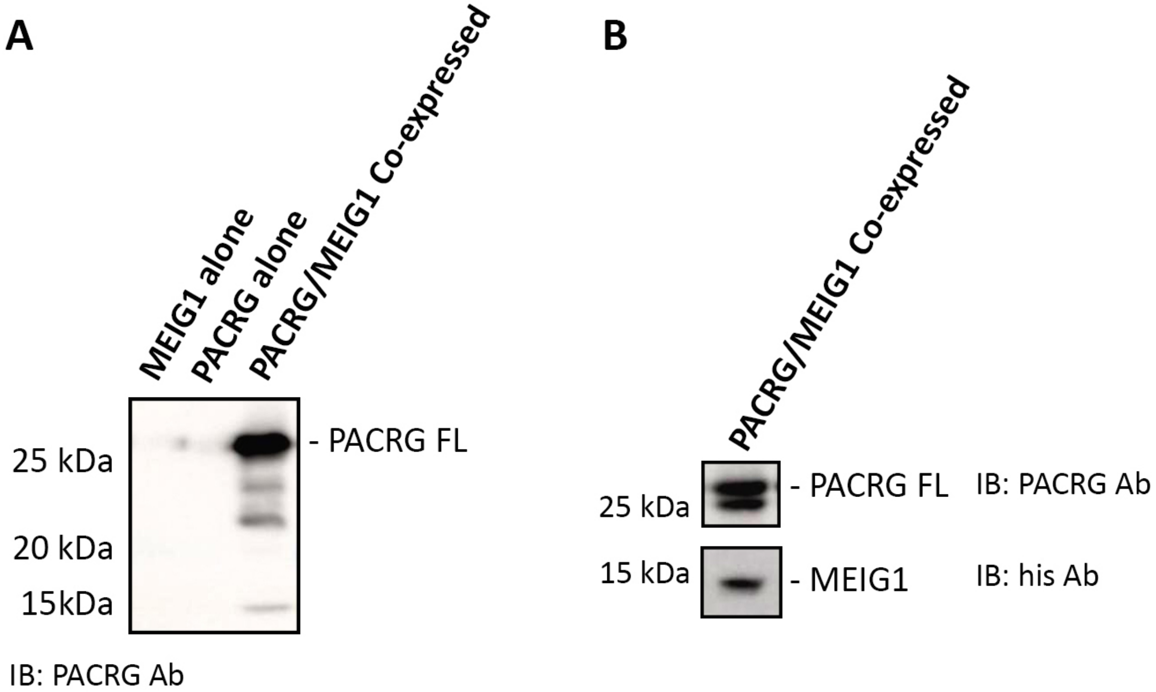


**Supplemental Figure 1. PACRG$_{FL}$:MEIG1 co-expression in *E. coli*. (A)** Cells expressing MEIG1 or PACRG alone showed little to no signal at 29 kDa, PACRG$_{FL}$ expected size, when blotted against PACRG. When PACRG was co-expressed with MEIG1, proteins levels significantly increased. Lysates were normalized by total protein concentration. **(B)** Cell lysates were blotted against PACRG and the his-tag on MEIG1 to indicate that both proteins were indeed expressed. Immunoblotting was performed with PBS with 0.1% Tween 20 with either PACRG mouse monoclonal antibody (1:2000, Santa Cruz Biotechnology, Inc.) or His-tag polyclonal rabbit antibody (1:2000, Cell Signaling Technology) in 15 mL of PBST with 2% bovine serum albumin (BSA), shaking over night at 4°C.

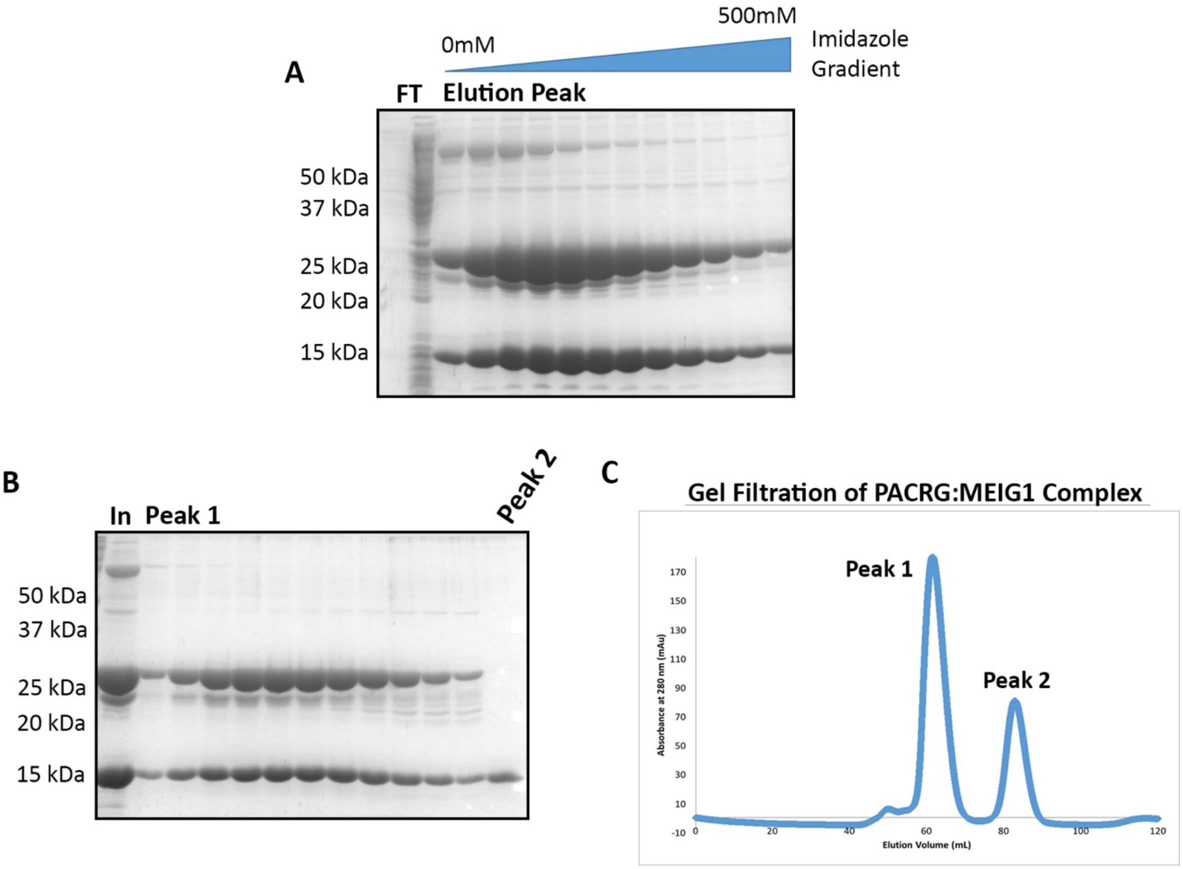


**Supplemental Figure 2**. **Purification of co-expressed PACRG$_{FL}$:MEIG1. (A)**. A richer media, TB, was used for E. coli growth and protein production followed by Ni-NTA gradient purification. The flow-through removed many impurities. An imidazole gradient (0-500 mM, blue bar) was used to separate the complex from *E. coli* proteins that bound to the column non-specifically. The complex was expressed in large amounts, PACRG$_{FL}$ at 29 kDa and MEIG1 at 16 kDa. The *E. coli* Lac Operon Repressor protein, large band above 50 kDa, was still present. **(B)**. Gel filtration was performed on the elutions from the Ni-NTA purification to remove large impurities like the Lac Operon Repressor. SDS-PAGE of gel filtration peaks. Peak1 corresponded to the complex, it was relatively pure enough to stop at this purification step. Peak 2 corresponded to free MEIG1 that did not bind to PACRG$_{FL}$. **(C)** Chromatogram of gel filtration, input was the elution fractions from Ni-NTA purification. The gel filtration separated the proteins successfully, the shoulder to the left of peak 1 corresponds to the Lac Operon Repressor. FT: flow through, In: input.

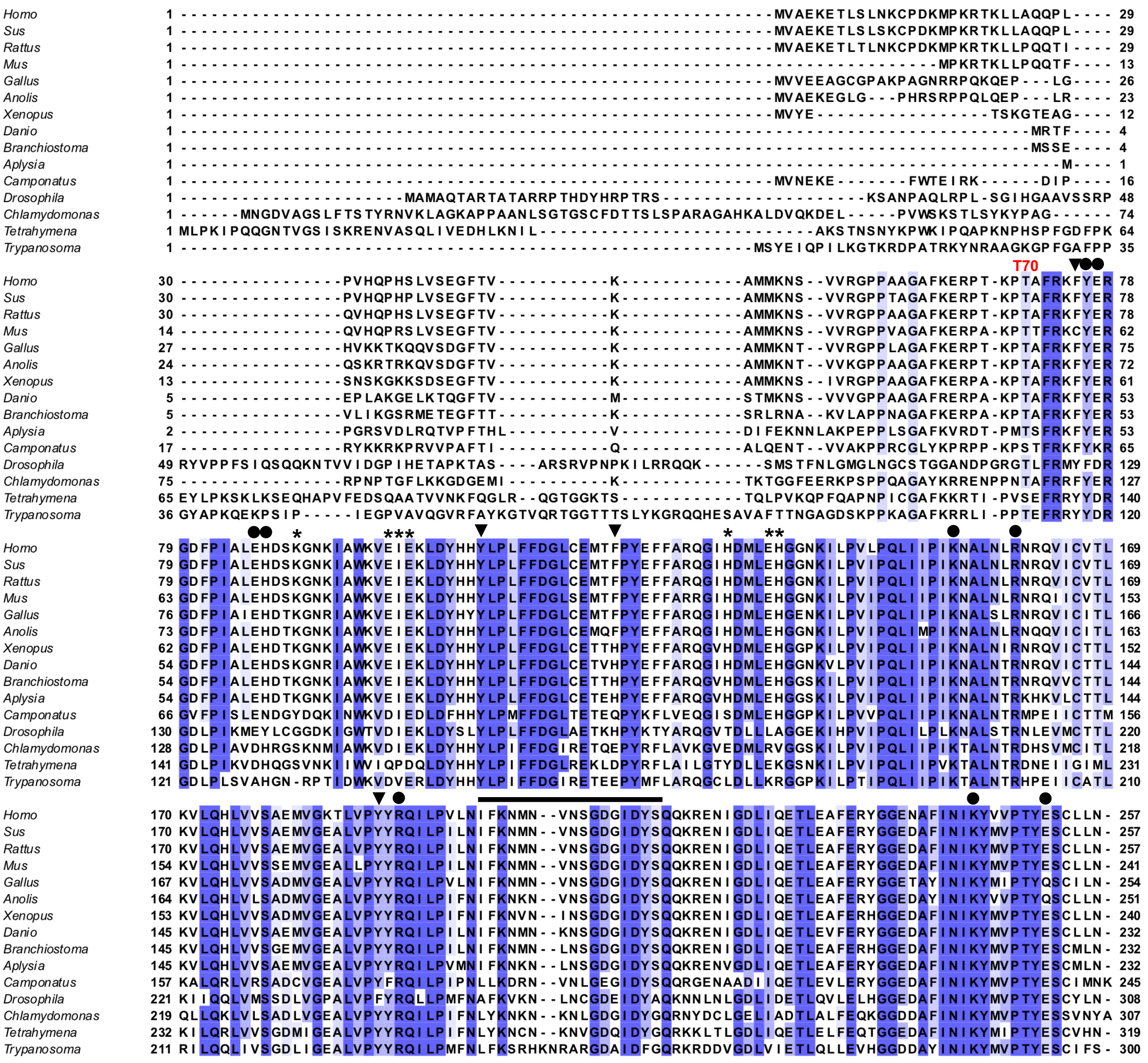


**Supplemental Figure S3**. **Sequence alignment of PACRG protein orthologs from different species.** PACRG N-terminal is highly variable across species, whereas the C-terminus is more conserved. Deletion constructs were made by deleting amino acids 1-69 (a.a. Thr70 labeled in red) in human PACRG. Positions labeled with black arrows indicate site where p-iodo-L-phenylalanine was incorporated for phasing. Asterisks indicate residues important for binding MEIG1. Black circles highlight residues that interact with tubulin in the axonemal doublet tubule structure.

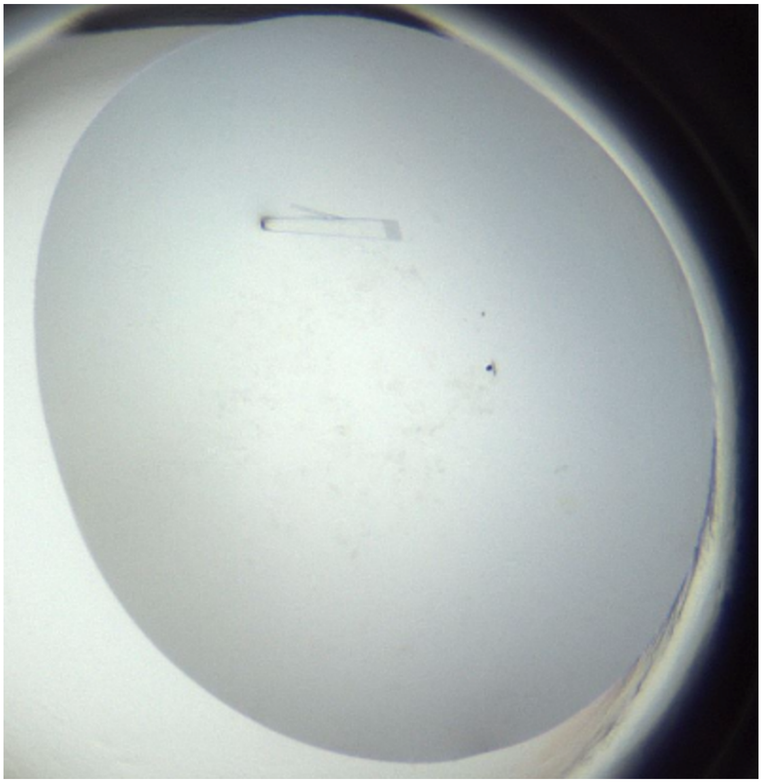

**Supplemental Figure S4**. **Crystal of the MEIG1:PACRG$^{\Delta 1\text{-}69}$ complex obtained by vapour diffusion.** The protein was crystallized at 1.7 mg/mL in 16 % w/v PEG 8000, 20 % v/v glycerol, 0.04 M $K_2HPO_4$.

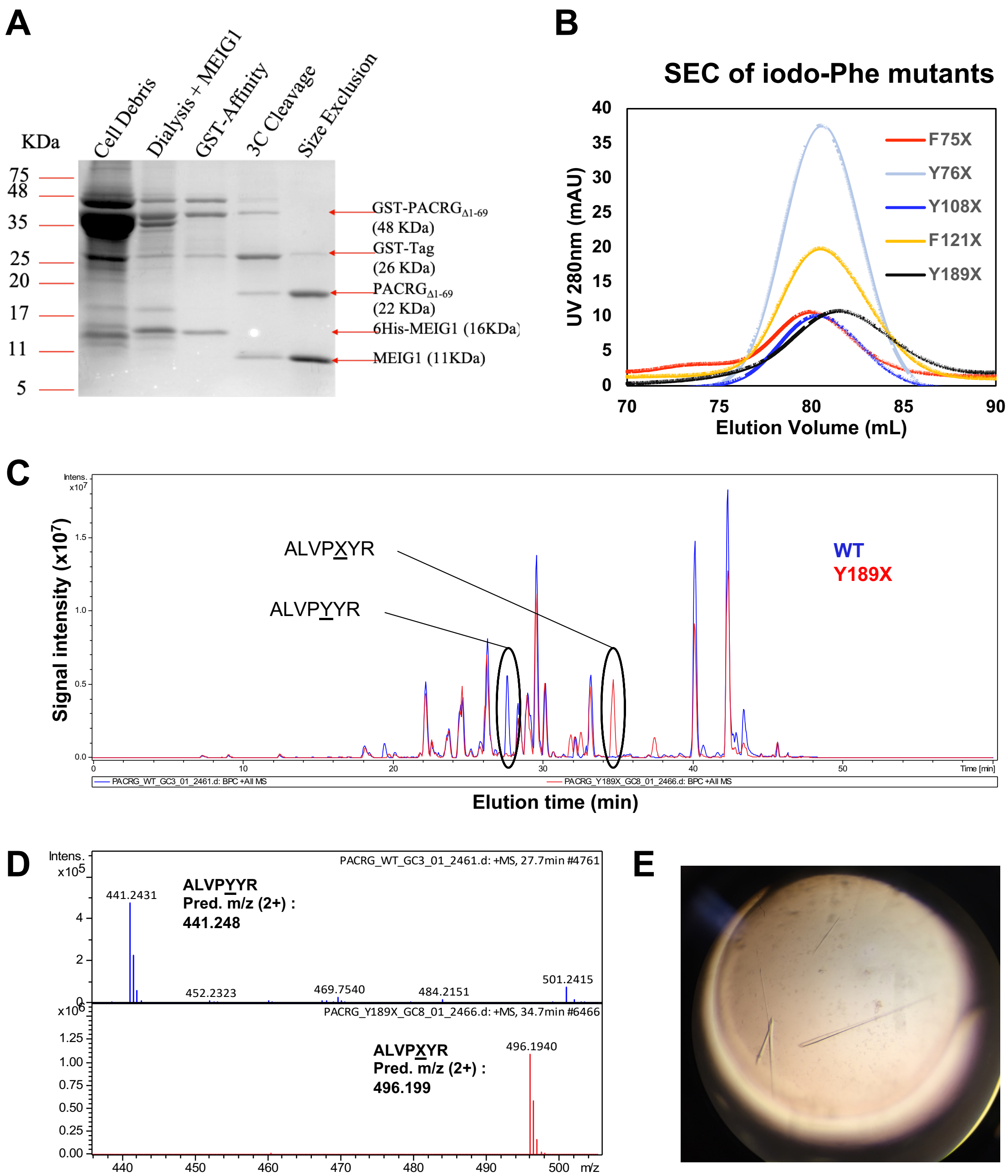


**Supplemental Figure S5**. **Purification of *p*-iodo-*L*-phenylalanine mutants of PACRG$^{\Delta 1\text{-}69}$ bound to MEIG1. (A)** Fractions from the Y189X PACRG $_{\Delta 1\text{-}69}$ mutant inclusion body purification migrated on SDS-PAGE gel and stained with Coomassie Brilliant Blue. The fractions consisted of protein from the cell debris collected after removing the cleared lysate (Cell Debris), after dialysis and addition of MEIG1 (Dialysis+MEIG1), after GST-affinity chromatography (GST-Affinity), following overnight 3C protease cleavage (3C cleavage), and after size-exclusion chromatography (Superdex 75). **(B).** UV 280 nm intensity for all eluted mutants on size-exclusion chromatography. **(C)** Extracted ion chromatograms of tryptic peptides from the wild type (WT) and Y189X mutant ofPACRG$^{\Delta 1\text{-}69}$ bound to MEIG1. All peptides are identical except the peptide spanning amino acids 185-191, which contains iodo-phenylalanine at position 189. **(D)** The precursor spectra indicate the mass shift induced by the incorporation of iodo-phenylalanine. **(E)** Crystals of the MEIG1:PACRG$^{\Delta 1\text{-}69}$ Y189X complex.

```
PACRG_short    1 MVAEKETLSLNKCPDKMPKRTKLLAQQPLPVHQPHSLVSEGFTVKAMMKNSVVRGPPAAGAFKER 65
PACRG_long     1 MVAEKETLSLNKCPDKMPKRTKLLAQQPLPVHQPHSLVSEGFTVKAMMKNSVVRGPPAAGAFKER 65

PACRG_short   66 PTKPTAFRKFYERGDFPIALEHDSKGNKIAWKVEIEKLDYHHYLPLFFDGLCEMTFPYEFFARQG 130
PACRG_long    66 PTKPTAFRKFYERGDFPIALEHDSKGNKIAWKVEIEKLDYHHYLPLFFDGLCEMTFPYEFFARQG 130

PACRG_short  131 IHDMLEHGGNKILPVLPQLIIPIKNALNLRNRQVICVTLKVLQHLVVSAEMVGKALVPYYRQILP 195
PACRG_long   131 IHDMLEHGGNKILPVLPQLIIPIKNALNLRNRQVICVTLKVLQHLVVSAEMVGKALVPYYRQILP 195

PACRG_short  196 VLNIFKNMN---------------------------------------VNSGDGIDYSQQKRENI 221
PACRG_long   196 VLNIFKNMNGSYSLPRLECSGAIMARCNLDHLGSSDPPTSASQVAEIIVNSGDGIDYSQQKRENI 260

PACRG_short  222 GDLIQETLEAFERYGGENAFINIKYVVPTYESCLLN                              257
PACRG_long   261 GDLIQETLEAFERYGGENAFINIKYVVPTYESCLLN                              296
```

**Supplemental Figure S6**. **Sequence alignment of PACRG isoforms.** The disordered loop in between α6 and α7 in the crystal structure of the short isoform of PACRG is highlighted with a blue box.

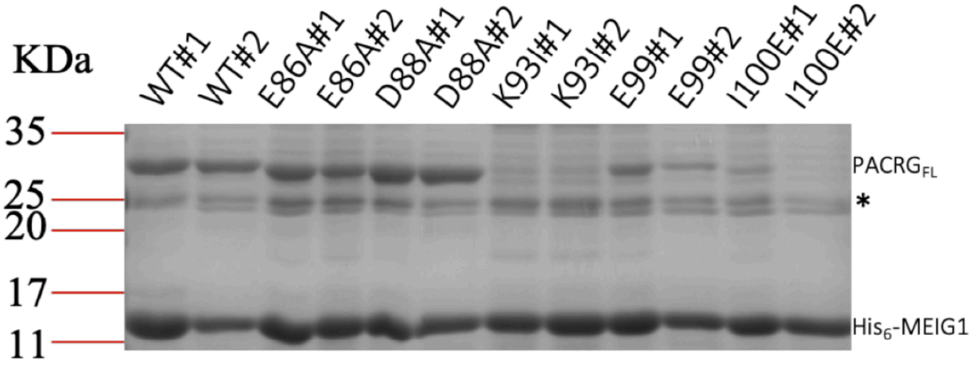


**Supplemental Figure S7**. **Interaction assay for the binding of PACRG mutants to His6-MEIG1.** WT or mutants PACRG$^{FL}$ were expressed with His$_6$-MEIG1 from a single pRSF-DUET plasmid. Following expression in E.coli, the cell lysate was incubated with Co-NTA affinity resin to pull-down on His$_6$-MEIG1. The eluate was loaded directly on SDS-PAGE and stained with Coomassie (example shown here). Densitometry was performed on both the PACRGFL and His$_6$-MEIG1 bands to calculate the ratios showed in Figure 2B. Each mutant was tested 4 times independently. The asterisk indicates a contaminant protein co-eluting non-specifically on Co-NTA.

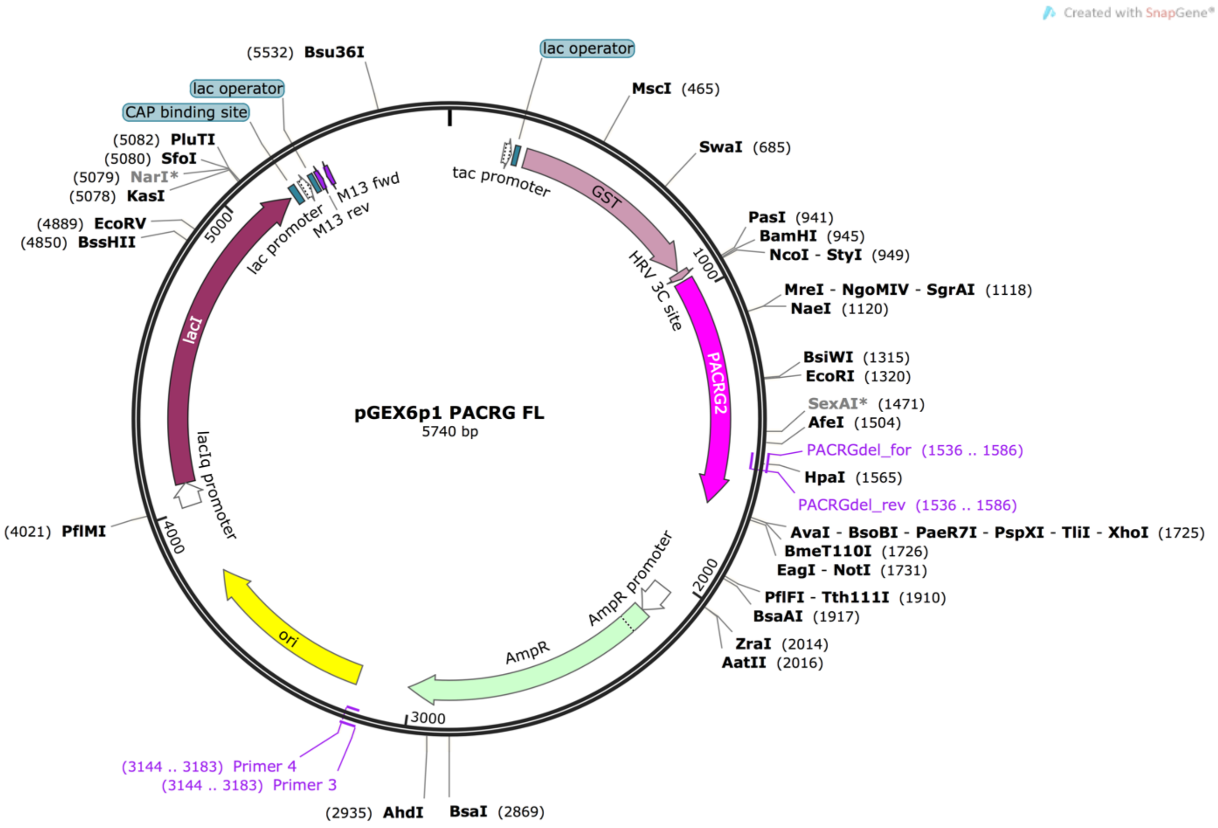


CC**ATG**GTTGCGGAAAAAGAAACCCTGTCTCTGAACAAATGCCCGGACAAAATGCCGAAAC
GTACCAAACTGCTGGCGCAGCAGCCGCTGCCGGTTCACCAGCCGCACTCTCTGGTTTCTG
AAGGTTTCACCGTTAAAGCGATGATGAAAAACTCTGTTGTTCGTGGTCCGCCGGCGGCGG
GTGCGTTCAAAGAACGTCCGACCAAACCGACCGCGTTCCGTAAATTCTACGAACGTGGTG
ACTTCCCGATCGCGCTGGAACACGACTCTAAAGGTAACAAAATCGCGTGGAAAGTTGAAAT
CGAAAAACTGGACTACCACCACTACCTGCCGCTGTTCTTCGACGGTCTGTGCGAAATGACC
TTCCCGTACGAATTCTTCGCGCGTCAGGGTATCCACGACATGCTGGAACACGGTGGTAAC
AAAATCCTGCCGGTTCTGCCGCAGCTGATCATCCCGATCAAAAACGCGCTGAACCTGCGT
AACCGTCAGGTTATCTGCGTTACCCTGAAAGTTCTGCAGCACCTGGTTGTTTCTGCGGAAA
TGGTTGGTAAAGCGCTGGTTCCGTACTACCGTCAGATCCTGCCGGTTCTGAACATCTTCAA
AAACATGAACGTTAACTCTGGTGACGGTATCGACTACTCTCAGCAGAAACGTGAAAACATC
GGTGACCTGATCCAGGAAACCCTGGAAGCGTTCGAACGTTACGGTGGTGAAAACGCGTTC
ATCAACATCAAATACGTTGTTCCGACCTACGAATCTTGCCTGCTGAAC**TAA**CTCGAG

**Supplemental Figure S8**. **Vector map of pGEX6p1-PACRG short isoform (257 a.a.)**. The sequence of the PACRG coding sequence (codon-optimized for E. coli expression), with the NcoI and XhoI restriction sites underlined, is shown below. The initiator (Met) and stop codons are in bold.

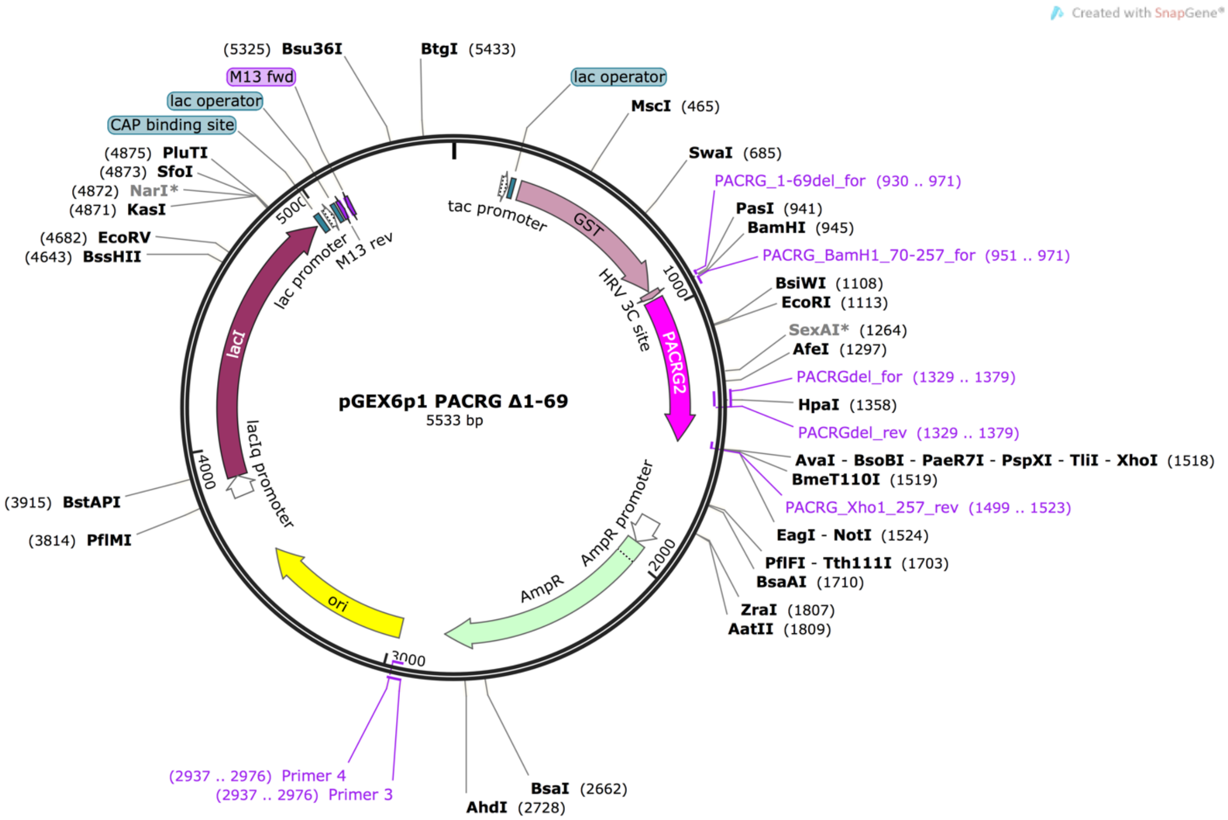


**Supplemental Figure S9. Vector map of pGEX6p1-PACRG$^{\Delta1\text{-}69}$.**

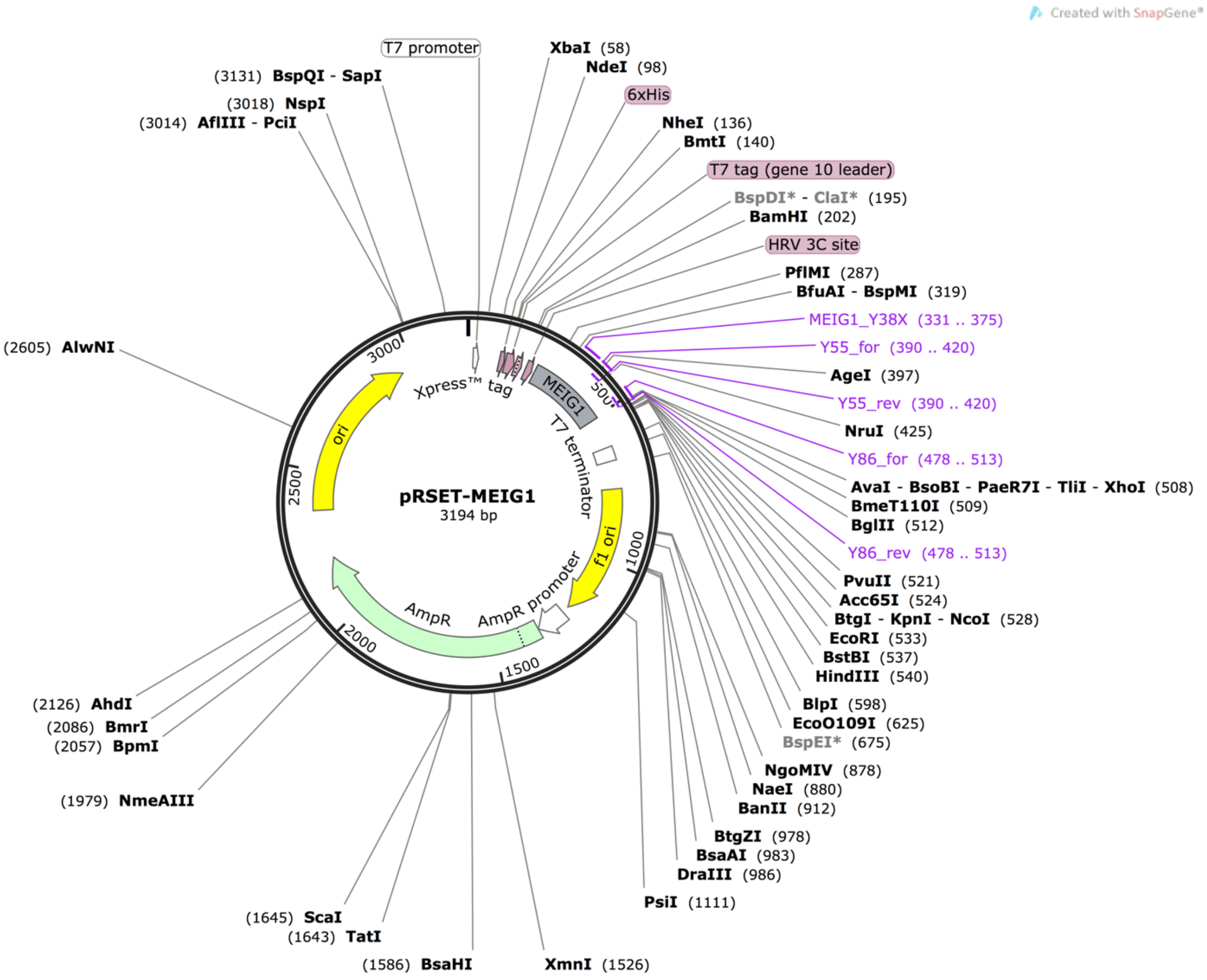


CAT**ATG**CGGGGTTCTCATCATCATCATCATCATGGTATGGCTAGCATGACTGGTGGACAGC
AAATGGGTCGGGATCTGTACGACGATGACGATAAGGATCGATGGGGATCCCTGGAAGTTC
TGTTTCAGGGTCCGCTGGGTAGCATGGCAAGCAGTGATGTTAAACCGAAAAGCGTTAGCC
ATGCCAAAAAATGGTCAGAAGAAATCGAAAACCTGTATCGTTTTCAGCAGGCAGGTTATCG
TGATGAAACCGAATATCGTCAGGTTAAACAGGTTAGCATGGTTGATCGTTGGCCTGAAACC
GGTTATGTTAAAAAACTGCAGCGTCGCGATAACACCTTCTACTATTACAATAAACAGCGCGA
GTGCGACGATAAAGAAGTGCATAAAGTTAAAATCTATGCCTAT**TAA**CTCGAG

**Supplemental Figure S10. Vector map of pRSET-His$_6$-MEIG1**. The sequence of the open-reading frame comprising the His$_6$-tag (red), Nde1 and Xho1 sites (underlined) and codon-optimized sequence of MEIG1 (blue) is shown below. The initiator (Met) and stop codons are in bold.

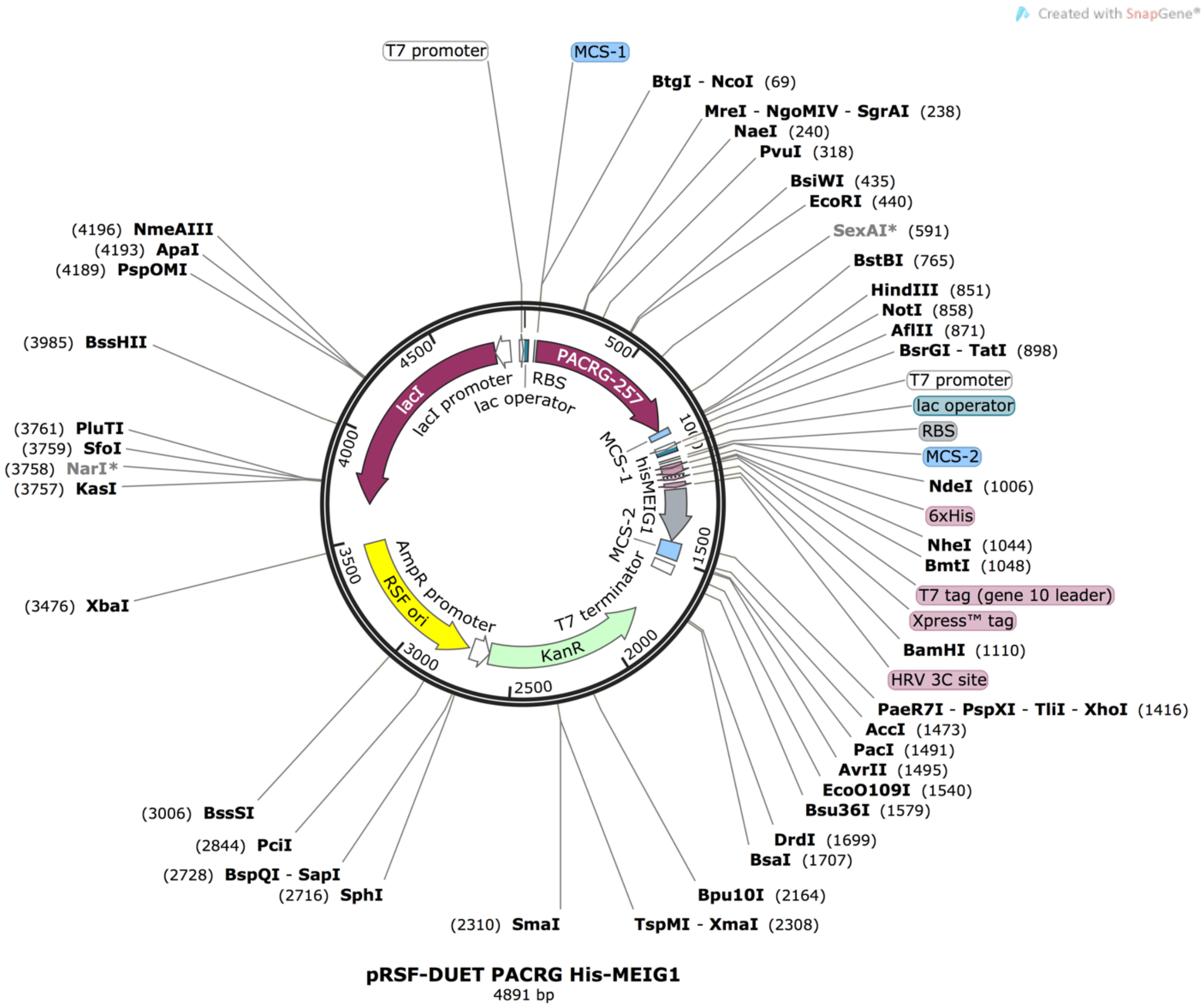


**Supplemental Figure S11. Vector map of pRSF-DUET PACRG$^{FL}$-His$_6$MEIG1.**
